# Haplotype-resolved genome assemblies and population resequencing reveal no evidence for genetic sex determination in the dioecious conifer Taxus baccata

**DOI:** 10.64898/2026.02.27.708407

**Authors:** Daniel Bross, Desanka Lazić, Malte Mader, Jannes Mittelbach, Martin Pippel, Stefan Kurtz, Emilia Pers-Kamczyc, Venket Raghavan, Hilke Schroeder, Laura Uelze, Sylke Winkler, Niels A. Müller, Birgit Kersten

## Abstract

Hundreds of plant lineages have independently evolved dioecy, the separation of female and male reproductive organs on different individuals. In all dioecious plants investigated at the molecular level to date, sex is determined genetically through a sex-determining region (SDR). However, SDRs have mostly been characterized in angiosperms, even though dioecy is far more common among gymnosperms. Here, we investigate sex determination in the dioecious conifer *Taxus baccata* L. (European yew). We generated chromosome-level haplotype-resolved genome assemblies, with a mean size of 10.04 Gb, for both haplotypes of one female and one male tree and resequenced 100 phenotypically sexed individuals at ∼20× coverage. Strikingly, reference-free *k*-mer analyses as well as reference-based coverage and association analyses failed to detect any signal of sex chromosomes or a sequence-differentiated SDR. These results thus leave two intriguing possibilities for sex determination in *T. baccata*, a genetic factor far more cryptic than any SDR described to date or a non-genetic mechanism.

## INTRODUCTION

Dioecious plants have evolved a variety of sex-determining systems^1,2^. These range from the well-known X/Y or Z/W systems, in which male-or female-heterogametic regions dominantly determine sex, to X/A balance systems, in which the dosage of the homogametic sex chromosome controls dioecy^3,4^. In plants, dioecy has evolved many times independently^5–7^. New genomic methods finally allow the identification and characterization of the underlying sex-determining regions (SDRs), that is, the genomic regions in which the sex-determining genes are located. This may reveal higher-level patterns and biological insights^8^. Notably, most studies on SDRs so far were conducted in angiosperms. While dioecy is present only in 6% of angiosperms^9^, it is far more common among gymnosperms, where about 65% of species are dioecious^5^. Within the gymnosperms, the SDRs have been studied at a molecular level in *Ginkgo biloba, Cycas panzhihuaensis* and *Welwitschia mirabilis*, with all three species exhibiting X/Y sex-determining systems^10–12^. Beyond this, several dioecious gymnosperms have been studied by karyotyping^13^.

The genus *Taxus* (Taxaceae) comprises 16 genetically defined lineages distributed across the Northern hemisphere^14^. *Taxus baccata* L. is the only species on the European continent^14^. All *Taxus* species except *T. canadensis*^15^ are dioecious. They are wind-pollinated understory trees or shrubs, with animal-dispersed seeds^16^. While cosexual individuals were reported in *T. brevifolia*, where female reproductive structures were found on 29.3% of predominantly male trees^16,17^, cosexuals appear to be exceedingly rare in most cases. In *T. cuspidata*, for example, no cosexual individuals were found in three studied populations^18^. For *T. baccata*, a fraction of 0.13% cosexual individuals was reported in seven Central European populations, in which singular branches of male trees formed female strobili in addition to male strobili^19^. Moreover, populations of *T. baccata* appear to generally have an even sex ratio^20^. These results indicate that most *Taxus* species in fact feature regular dioecy. While *Taxus* has received attention from numerous researchers in recent years, mainly due to its capacity to produce the anticancer drug paclitaxel^21,22^, the sex-determining systems in this genus remain elusive. The publication of a reference genome for *T. wallichiana* proposed a Z/W system with a putative SDR located on chromosome 12 (ref.^23^). However, this suggestion was based on indirect evidence, specifically length differences between the two reference haplotypes. For *T. baccata*, a karyotype analysis showed the absence of heteromorphic sex chromosomes^24^. So far, no sex-specific markers have been found^25^.

In this study, we aimed to elucidate the sex-determining system of *T. baccata* by generating high-quality long-read (PacBio HiFi) sequencing data and chromatin conformation (Hi-C) data to assemble chromosome-level haplotype-resolved genome assemblies for both haplotypes of a female and a male individual. Additionally, we generated short-read resequencing data from 100 individuals of known sex to analyze potential sex-specific sequences. Strikingly, our analyses demonstrate the absence of any evidence for sex-linked sequences in different reference-free and reference-based analyses, indicating that sex determination may be controlled by a cryptic genetic factor or a non-genetic molecular mechanism in *T. baccata*.

## RESULTS

### Taxus baccata genome assemblies

To construct female and male chromosome-level haplotype-resolved genome assemblies of *T. baccata*, we generated PacBio HiFi long-read sequencing data with a coverage of ∼36× and ∼41× (359.32 and 410.02 Gb), and short reads from Hi-C libraries with a coverage of ∼63× and ∼50× (632.11 and 499.10 Gb) for a female and a male individual, respectively.

The total sizes of the four final haplotype assemblies range from 9.87 to 10.19 Gb. Merqury^26^ *k*-mer completeness values were 99.3% and 98.6% for the combined female and male haplotypes, respectively, or from 75.7% to 76.6% for the individual assemblies (QVs 65.6 – 66.2) (Table 1, Supplementary Table 1). In each assembly, 12 chromosomes contain 99.6% to 100% of the haplotype-assigned bases, and N50 values range from 12.2 to 13.6 Mb for contigs and from 903.6 to 928.7 Mb for scaffolds. The four assemblies contain between 95.6% and 96.2% of embryophyte BUSCOs (embryophyta_odb12) (Table 1, Supplementary Table 1), indicating a relatively high completeness. The GC content ranged from 36.58% to 36.70% (Table 1, Supplementary Table 1). Telomeric repeats were identified in the terminal regions of 40 out of the 48 chromosomes across the four assemblies. The Hi-C heatmaps of the four assemblies show the successful scaffolding into chromosomes (Extended Data Fig. 1).

**Table 1.** Overview of the four genome assemblies and their annotation.

| **Assembly** | **Tb_female-h1** | **Tb_female-h2** | **Tb_male-h1** | **Tb_male-h2** |
|---|---|---|---|---|
| Genome size [Gb] | 10.19 | 10.11 | 9.87 | 9.98 |
| GC content [%] | 36.7 | 36.7 | 36.6 | 36.6 |
| Number of scaffolds | 408 | 266 | 100 | 87 |
| Number of gaps | 1667 | 1593 | 1558 | 1586 |
| Scaffold N50 [Mb] | 928.7 | 921.3 | 903.6 | 924.8 |
| Number of contigs | 2075 | 1859 | 1658 | 1673 |
| Contig N50 [Mb] | 12.2 | 12.2 | 13.6 | 13.6 |
| Contig L50 | 242 | 237 | 210 | 219 |
| Anchored [%] | 99.6 | 99.8 | 100 | 100 |
| Merqury *k*-mer completeness [%] | 76.6 | 76.4 | 75.7 | 76.5 |
| Merqury QV | 65.6 | 65.6 | 66.2 | 66.2 |
| Combined Merqury *k*-mer completeness [%] | 99.3 |  | 98.6 |  |
| Combined Merqury QV | 65.6 |  | 66.2 |  |
| Complete (C) BUSCOs [%] | 95.9 | 96.2 | 95.6 | 95.8 |
| Single-copy (S) [%] | 90.2 | 90.5 | 89.8 | 90.1 |
| Duplicate (D) [%] | 5.7 | 5.7 | 5.8 | 5.7 |
| Fragmented (F) [%] | 1.8 | 1.6 | 1.4 | 1.9 |
| Missing (M) [%] | 2.3 | 2.2 | 3.0 | 2.3 |
| Complete with internal stop codon (E) [%] | 17.9 | 18.0 | 18.6 | 17.0 |
| Total gene count | 35,060 | 39,283 | 33,907 | 36,096 |
| Total transcript count | 44,597 | 48,657 | 42,568 | 45,383 |
| Mono:multi-exonic transcript ratio | 0.48 | 0.67 | 0.46 | 0.52 |
| Functional annotations [%] | 80.3 | 75.0 | 82.0 | 79.7 |

The four haplotype assemblies exhibit considerable structural variation (Fig. 1a, Extended Data Figs. 2-3, Supplementary Fig. 1). Syntenic regions cover between 85.5% and 86.9% of the genome sequences for all pairs of genome assemblies. The largest inversion spans almost 100 Mb on chromosome 8 and is supported by highly resolved Hi-C data (Extended Data Fig. 4) and long read mapping (Supplementary Fig. 2). This inversion is heterozygous in both reference individuals (Extended Data Figs. 2-4) suggesting that it may not be involved in sex determination. However, the assemblies also highlight large inversions that are heterozygous only in one of the two individuals, such as a ∼46 Mb inversion in the middle of chromosome 3 in the female individual (Extended Data Figs. 2-3, 5), which might be expected in the case of a female-heterogametic Z/W system (although our analyses below show no indication for such a system).

**Fig. 1:**
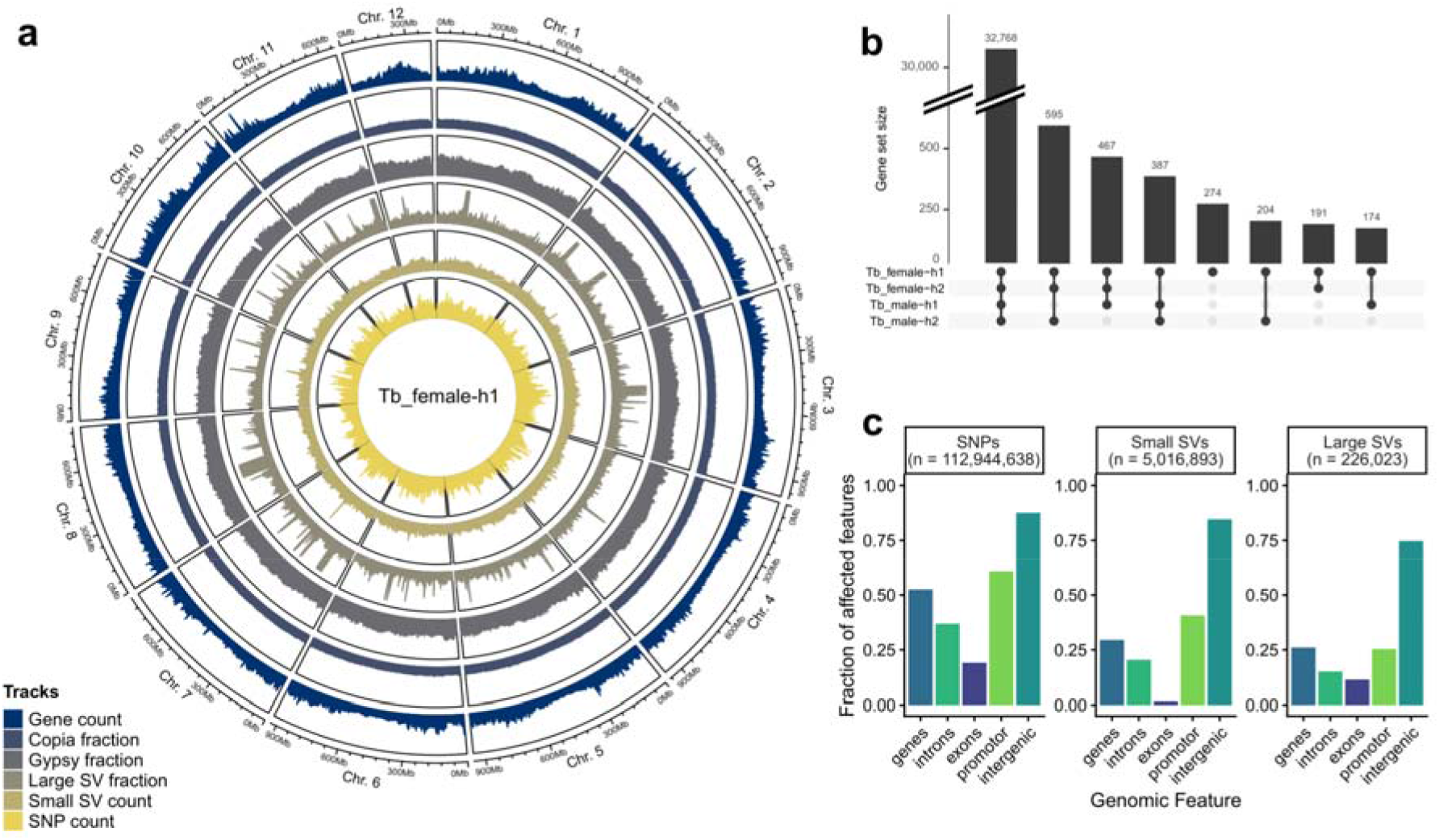
The assembly of female haplotype 1 reveals high levels of genomic variation and haplotype-specific genes. **a,** Circos plot of the Tb_female-h1 assembly. Statistics were calculated in 5 Mb sliding windows with 2.5 Mb steps. The following feature types are presented: Genes (counts per window); Copia elements, Gypsy elements, and large structural variants (> 50 bp) (fraction of the window covered by the respective feature type); small structural variants (<= 50 bp) and SNPs (counts per window). **b,** Comparisons of the gene sequences annotated in the Tb_female-h1 assembly with the other assemblies’ sequences, as determined by Liftoff, reveal presence-absence variation of genes. Each bar refers to one combination indicated below the bar, with black dots marking presence and gray dots absence from the assembly. **c,** Fractions of genomic features that overlap with different types of variants, i.e., SNPs, and small (<= 50 bp) or large structural variants (> 50 bp) reported by SyRI show conservation of genes, and specifically exons compared to intergenic regions. The regions up to 5 kb upstream of a gene are considered promoter regions.

Using both short-read and long-read RNA sequencing data, we generated structural annotations for each of the four haplotypes. Across the four assemblies, annotations with BRAKER3 yielded between 33,907 and 39,283 gene models (Table 1, Supplementary Table 1), excluding transposable elements, which were soft-masked prior to annotation. Notably, this soft-masking step covered between 82.78% and 83.80% of the assembly sequences, highlighting the extensive presence of transposable elements and low-complexity regions in the assemblies. The ratio of monoexonic to multiexonic gene models varied between 0.46 and 0.67 in the four assemblies. Functional annotations could be assigned to 74.98% to 82.04% of the transcripts in these gene models. While the gene density in the centromeric regions decreases as expected, the density of the repeat elements appears to be evenly distributed across the chromosomes (Fig. 1a, Supplementary Fig. 1). Using the machine learning approach Helixer as a complementary annotation method, we identified slightly higher numbers of gene models ranging from 40,504 to 41,303.

By transferring the sequences corresponding to the BRAKER3-gene annotations of the female haplotype 1 assembly to the other three assemblies using Liftoff, we identified genes shared among the four haplotypes, with respect to the assemblies’ sequences and not the individual annotations, but also genes specific to certain assemblies (Fig. 1b). We applied the same approach to each of the other three haplotypes (Supplementary Fig. 3). As expected, the majority of genes were identified in all four haplotypes. Nevertheless, more than 2,000 genes were missing in at least one haplotype, highlighting the potential relevance of gene presence-absence variation. A list of all haplotype-specific genes is given in Supplementary Table 2. Gene Ontology (GO) terms overrepresented among the haplotype-specific genes varied across the four assemblies and did not point to haplotype-specific gene clusters involved in sex determination, as might be expected in the case of an extended W or Y chromosome-specific hemizygous region (Supplementary Fig. 4). In addition to the gene presence-absence variants, there is a remarkable number of single nucleotide polymorphisms (SNPs) and structural variants (SVs) within our four assemblies. As expected, SVs are depleted in genes compared to SNPs (Fig. 1c, Supplementary Fig. 5).

### Population resequencing of Taxus baccata

To search for an SDR, we resequenced 103 samples for which the phenotypic sex was assessed before sample collection. From these samples (Supplementary Table 3), species identity was confirmed with molecular markers^27^ for 101, while two samples were identified as *T.* × *media* hybrids and excluded from further analysis. One additional sample was excluded due to a cosexual branch detected in the year after sampling. This resulted in a final set of 50 female and 50 male *T. baccata* samples, with 62 samples originating from seven putatively autochthonous populations and 38 samples from botanical gardens and other sources (Fig. 2a). A microsatellite analysis confirmed that no samples were clonal duplicates prior to sequencing (Supplementary Table 4).

**Fig. 2:**
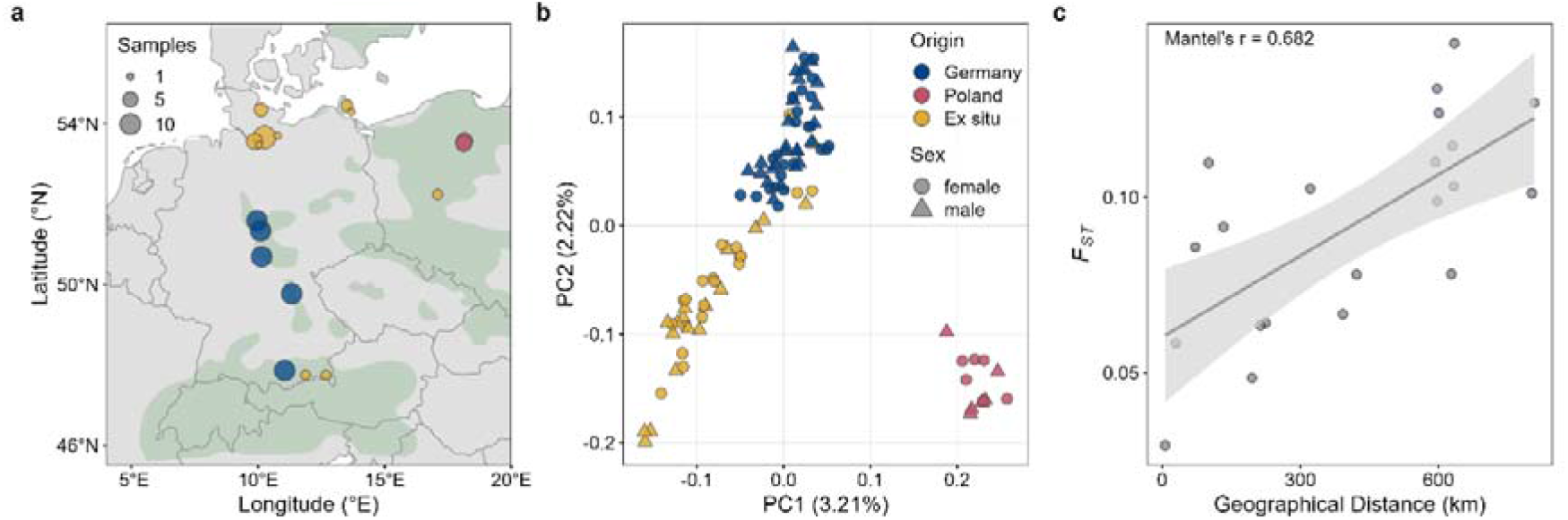
Genomic variation shows population structure in Taxus baccata across the sampling locations. **a,** Map of the collection sites of the 100 *T. baccata* samples. Autochthonous populations are marked with blue (Germany) or red points (Poland). Ex situ provenances include all samples from non-autochthonous populations and are marked with yellow points. Shapefile^28^ was used for the distribution of *T. baccata* individuals on the map. **b,** Principal component analysis based on a random subset of genome-wide SNPs called from mappings to Tb_female-h1 (Material and Methods) shows population structure among the 50 female (circle) and 50 male (triangle) *T. baccata* individuals. Each point depicts one individual. Colors indicate origin as in (a). **c,** Pairwise *F*_*ST*_ estimates of the seven autochthonous populations show relatively high genetic differentiation and significant isolation by distance (Mantel’s *r* = 0.682, *p* = 0.027).

Mapping the short-read resequencing data of the 100 samples to the four haplotype assemblies showed an average coverage of 20.8× per sample. Variant calling detected between 969 million and 971 million SNPs before filtering. The population structure of the dataset was explored by selecting a random subset of SNPs and performing a principal component analysis (PCA) (Fig. 2b, Supplementary Fig. 6). In general, samples from autochthonous populations in Germany formed partially overlapping clusters adjacent to (or overlapping with) the samples of ex situ provenances from botanical gardens and other sources, while samples originating from Poland clustered separately (Fig. 2b). Importantly, female and male individuals were homogeneously distributed among clusters, indicating that sex is not confounded by population structure in our dataset. Genetic differentiation (i.e., *F_ST_*) between pairs of populations was relatively high even among populations only separated by a few hundred kilometers (Fig. 2c). In general, genetic differentiation highlighted isolation by distance (Mantel’s *r* = 0.682, *p* = 0.027), with distant populations being more differentiated than adjacent ones.

### Reference-free k-mer analyses provide no evidence for W- or Y-chromosomal sequences

To determine the heterogametic sex in *T. baccata*, we first applied a reference-free approach by analyzing *k*Umers that are enriched in (or specific to) one sex, termed putative female-specific *k*-mers (FSKs) and putative male-specific *k*-mers (MSKs), based on the short reads from the 100 sexed individuals described above. We first compared *k*-mers from all 50 female and all 50 male samples under different criteria for sex-specificity. Specifically, we searched for *k-*mers that are present in a minimum fraction of samples in the target group (e.g., female samples for FSKs) and absent in at least the same fraction of samples in the non-target group (i.e., male samples for FSKs). This analysis resulted in similar counts of FSKs and MSKs under relaxed filter criteria, i.e. 55-70% sex-specificity, but none at filters stricter than 70% (Fig. 3a). When we split the samples into ten sets of randomly chosen groups of five females and five males, or into the different autochthonous populations, and tested the abundance of completely sex-specific *k*-mers (corresponding to a 100% filter in Fig. 3a) in each set, we found no consistently elevated counts of FSKs or MSKs (Fig. 3b; Supplementary Fig. 7). Notably, none of the *k*-mers identified as sex-specific in 5 vs. 5 comparisons occurred in more than 3 sets (Fig. 3c). The results of the same analyses looked strikingly different when repeated with short-read resequencing data of the dioecious gymnosperm *Ginkgo biloba*, a species with a similar genome size but a known X/Y-system of sex determination^10,29^. Here, the total sample set of 50 female and 50 male samples exhibited the expected male-biased signal especially under strict filtering criteria (Fig. 3d). Interestingly, the sets of five female and five male samples did not consistently show elevated counts of MSKs compared to FSKs (Fig. 3e). Instead, the ratios appeared to be driven by randomly differentiated genomic regions pointing to a relevant difference of species with very large genomes compared to species with smaller genomes^30,31^. However, the MSK signal became unambiguous when at least 5 sets (i.e., 25 vs. 25 individuals) were combined, and multiple *k*-mers were consistently identified as MSKs across all ten sets (Fig. 3f).

**Fig. 3:**
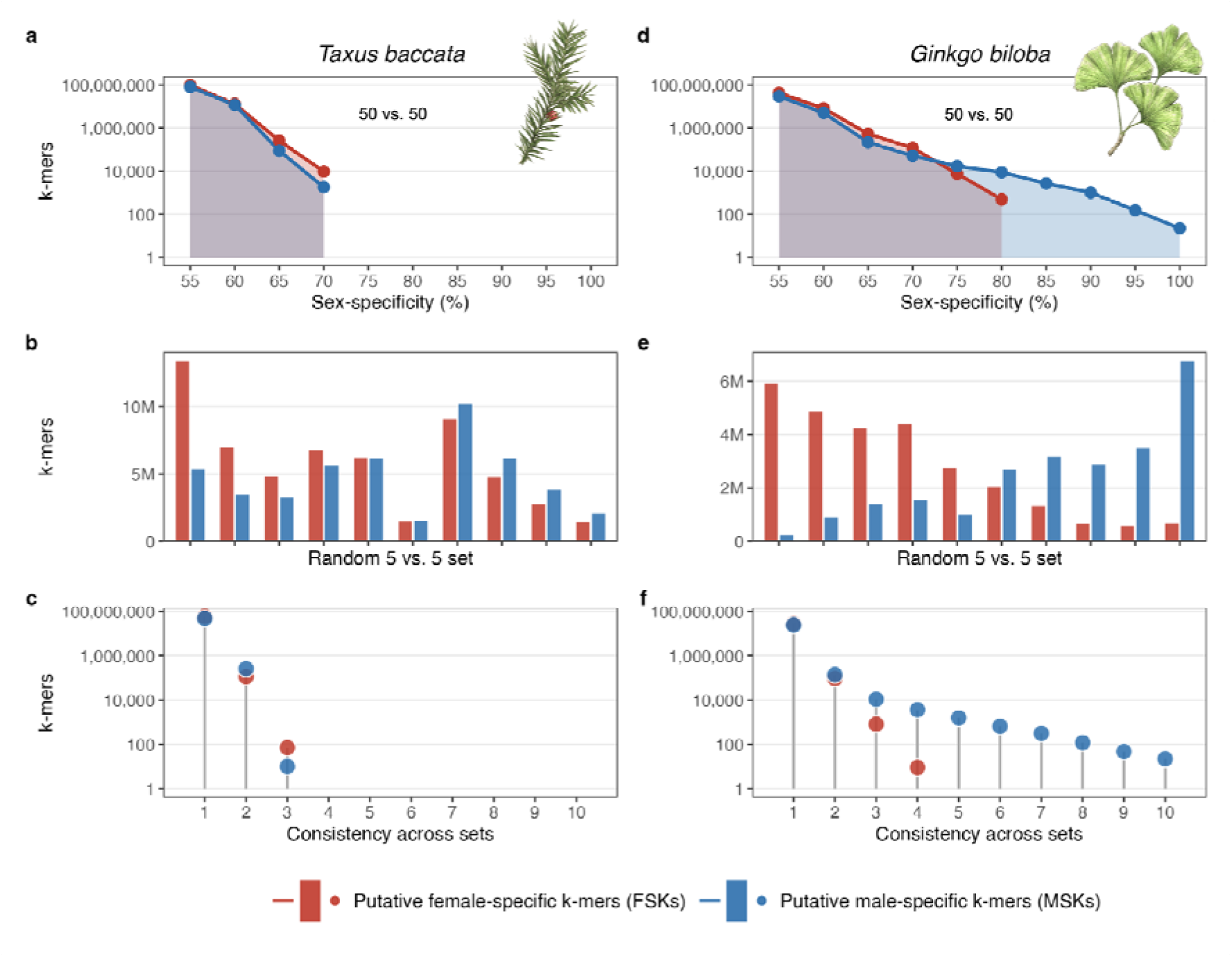
Reference-free k-mer analyses in Taxus baccata and Ginkgo biloba provide no evidence of sex-specific sequences in *T. baccata*. **a,d** Counts of putative female-specific k-mers (*FSKs*) and putative male-specific k-mers (*MSKs*) for different filtering criteria in sets of 50 female and 50 male samples of T. baccata (a) or G. biloba (d). The sex-specificity corresponds to the minimum fraction of target samples containing the *k-mer* while the same fraction of non-target samples does not contain that k-mer. Photos of twigs depict the two species. **b,e** Counts of putative sex-specific k-mers in 10 random sets of 5 female and 5 male samples of *T. baccata* (b) and G. biloba (e) demonstrate a high degree of noise in both species. Sets are ordered based on their FSK/MSK ratio. **c,f** Consistency of FSKs and MSKs across the sets shown in b,e. Consistency refers to the number of sets that contain a given FSK or MSK.

These analyses show that in dioecious species with large genomes, comparisons of small groups of 5 females vs. 5 males result almost entirely in random noise. The picture gets even worse when allowing one deviant individual per group (Supplementary Fig. 8). Segregating presence-absence variants, which appear widespread based on an analysis of a pangenome graph generated from our four T. baccata assemblies (Supplementary Fig. 9), can generate abundant group-specific sequences purely by chance. To push the random signal substantially below the truly sex-specific signal appears to require at least 25 individuals per sex in G. biloba (Fig. 3f). In larger groups of 50 vs. 50, a specificity threshold of 85% (i.e. k-mers present in at least 85% of one group and absent in 85% of the other group) is able to highlight genuinely sex-specific sequences (Fig. 3d). In contrast, the results for Taxus show no evidence whatsoever for specific W- or Y-chromosomal sequences. Could this indicate the absence of a genetic system of sex determination? Or may a potentially small SDR with relatively high levels of neutral differentiation, consistent with the *F*_*ST*_ estimates from above, avoid sex-specific *k*-mers?

### Reference-based analyses across haplotypes fail to identify a sex-determining region

To explore the possibility of an SDR with high differentiation within groups of females or males, we performed several reference-based analyses. If in fact sex-specific *k*-mers were masked due to secondary non-sex-linked mutations, we may still expect typical signatures of SDRs such as increased genetic differentiation or differences in nucleotide diversity between groups of female and male individuals. However, neither genetic differentiation (*F_ST_*) nor nucleotide diversity (π) showed any sex-specific patterns as expected from an SDR (Fig. 4a,b,h,i; Extended Data Fig. 6). Only the centromeric region of chromosome 6 appears to feature two low-recombination haplotype blocks that are unevenly distributed across our groups of females and males, as seen by slightly increased genetic differentiation. However, segregation patterns of homozygous and heterozygous genotypes, separating the two haplotype blocks along this region, are not consistent with an SDR (Supplementary Fig. 10). There are thus no indications of any extended SDR in the *T. baccata* genome from reference-based analyses either. Still, these analyses cannot rule out a small SDR or even less so a single causal mutation, controlling the activity of a sex determination factor. Such a mutation should, however, be identified as a sex-associated variant in a genome-wide association study (GWAS).

**Fig. 4:**
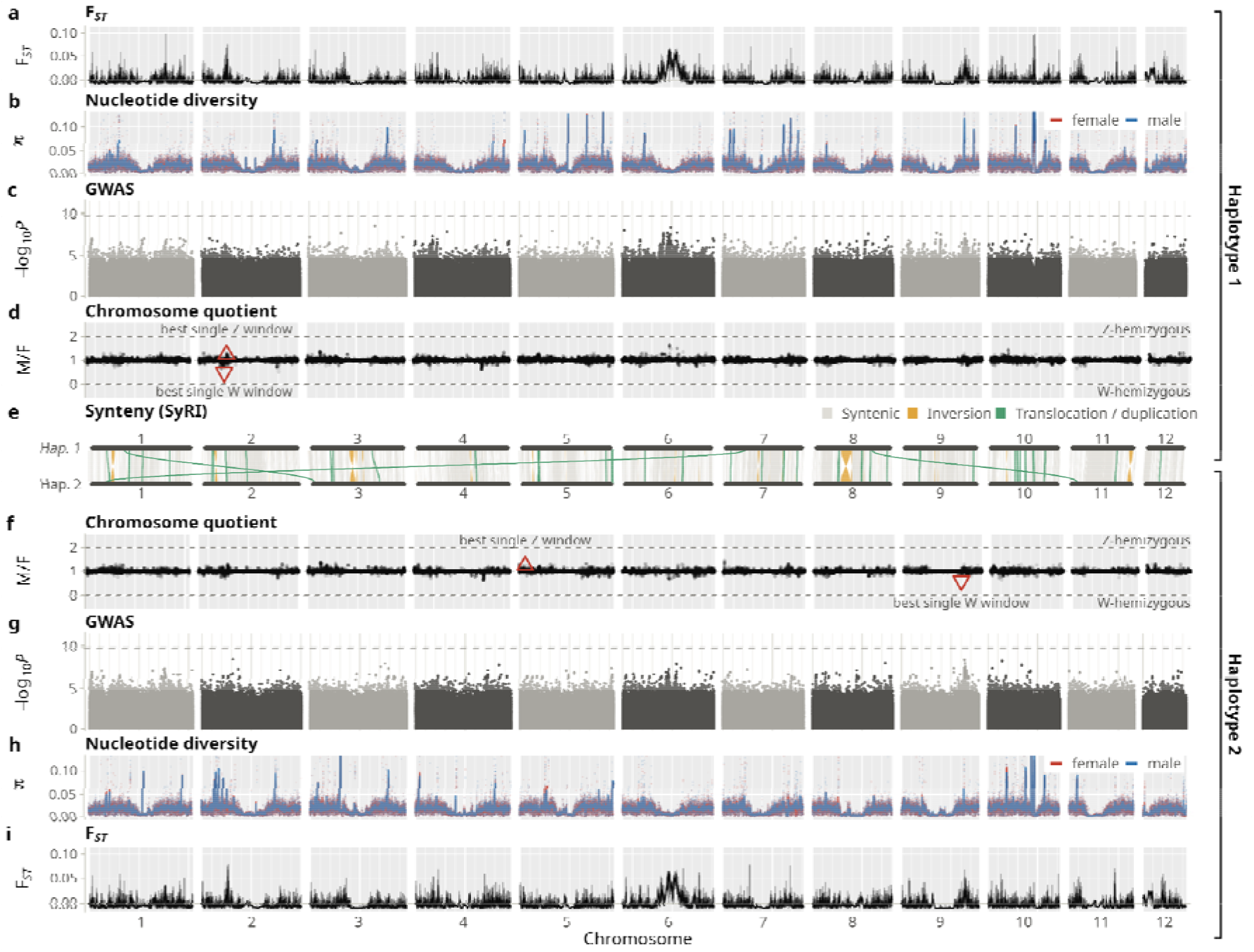
Reference-based analyses of 100 individuals across the two female reference haplotypes fail to identify a sex-determining region in Taxus baccata. **a,i** Genetic differentiation (F_ST_) between females and males was calculated in 100 bp windows and visualized as 100 kb averages. The black line shows the average across each plotting bin of about 2.4 Mb, the shaded band the range. **b,h** Nucleotide diversity π for females (red) and males (blue) in 100 kb windows is shown as points, with solid blue and red lines marking a sliding average across 5 Mb regions. **c,g** Manhattan plots show GWAS results (i.e., -log10 P-values) for sex of ca. 320 million biallelic SNPs. Horizontal dashed lines indicate 5% Bonferroni significance cutoff. **d,f** The chromosome quotient was calculated as the ratio of mean male to mean female (M/F) short-read DNA sequencing coverage in blocks of 1,000 consecutive 100 bp windows (∼100 kb), normalized by the genome-wide median depth of each sample. Only windows with a normalized depth of at least 0.2 in at least 25 of the 50 females were kept. Single best Z (ratio closest to Z-hemizygosity of 2) and W (ratio closest to W-hemizygosity of 0) 100 bp windows are shown as red triangles. e SyRI was used to assess syntenic regions (gray), inversions (orange) and translocations/duplications (green) across the 12 chromosomes. The upper half of the figure depicts female haplotype 1 (Tb_female-h1), the lower half female haplotype 2 (Tb_female-h2), which represent the two haplotypes of the female reference assemblies. The 12 chromosomes are shown on the x-axis. All analyses used 100 resequenced individuals, split into females (n=50) and males (n=50).

We therefore performed genome-wide association studies based on the SNPs called from the short-read mappings against the four genome assemblies. These GWAS showed no significant SNPs against any of the four reference assemblies (Fig. 4c,g; Extended Data Fig. 6). The Q–Q plots confirmed the absence of SNPs deviating from the uniform *P*-value distribution expected by chance (Supplementary Fig. 11). Since short-read mapping is highly reliable in coding regions or across other non-repetitive functional elements, these results argue against a single causal SNP in a sex determination factor. But what if the causal sequence is hemizygous and thus not present in a regular GWAS (because of missing data), and again with too many secondary mutations to be picked up by the *k*-mer-based methods? To test for such sex-linked hemizygous sequences, we screened the four genome assemblies for hemizygous regions only present in females (W-hemizygosity) or males (Y-hemizygosity) across approximately 100 million windows of 100 bp each. Specifically, we assessed the “chromosome quotient”, that is the ratio of homogametic sex coverage to heterogametic sex coverage, which should be 2 in the case of a Z- or X-hemizygous and 0 in the case of a W- or Y-hemizygous region. However, the best single 100 bp windows did not even come close to these expected values (Fig. 4d,f; Extended Data Fig. 6). Also, the best Z and the best W window of the two female assemblies did not coincide, as would be expected if one of those carried a Z and the other one a W chromosome. None of the prominent inversions between haplotypes, as for example the one on chromosome 3 mentioned above (Extended Data Fig. 5), showed any signs of sex-linkage (Fig. 4e). The results of the reference-based analyses were similar for the two male assemblies without any evidence for possible X- and Y-chromosomal sequences (Extended Data Fig. 6). Together, these reference-based and reference-free analyses indicate that sex might not be determined genetically in *T. baccata*.

### A MADS-box gene shows male-specific expression and may act as a downstream factor of sex determination

Given the generally stable sex expression in T. baccata, it seems likely that sex is determined only once in an individual’s lifetime. To identify potential sex-determining factors expressed only in one sex and not the other, we performed an extensive RNA sequencing experiment. Specifically, we sequenced the total RNA of reproductive buds at an early stage of their development from 12 (or 11) female and 11 (or 10) male trees at two time points (i.e., June and July) and analyzed differential gene expression against all four genome assemblies (Figs. 5a-b, Supplementary Tables 5-12). If a primary sex-determining gene does not exhibit any sex-linked variant in its coding sequence, its sex-determining function must arise from differences in cis between its gametologs, either genetic or epigenetic. When these differences act at the level of transcription, we would expect largely monoallelic expression in the heterogametic sex. Using female haplotype 1, our differential expression analyses identified only two switch-like genes, *AGL11* being male- and Pyrophosphatase 4 being female-specific (Fig. 5c-f). While both genes were identified at both time-points, only the MADS-box gene *AGL11* was identified in all four assemblies making it a strong candidate for being involved in the molecular mechanism controlling sex expression in *T. baccata* (Supplementary Tables 5-12). Additionally, there were two other MADS-box genes (i.e., *MADS23*-1 and *MADS23*-2) that were male-biased in all analyses (Fig. 5a,b; Supplementary Tables 5-12), without being exclusively expressed in one sex only and thus potentially consistent with a Z/A balance system (Fig. 5e,f). In all cases, Y- and W-specific or Z-limited genes would exhibit monoallelic expression in the heterogametic sex. To test this for the four candidate sex-determining genes, we extracted heterozygous SNPs from the DNA sequencing data within the coding regions and assessed their allele balance in the mapping of the RNA sequencing data. Analysis of four to eight SNPs in several individuals and across three to four exons indicated that all genes exhibited expression from both alleles (Fig. 5g-j). It should be noted, however, that not every individual could be scored (Supplementary Fig. 12). Nevertheless, this makes their role as primary sex-determining factors unlikely. Rather, our results suggest that those genes might be important downstream targets in the molecular pathway regulating differential sex expression.

**Fig. 5:**
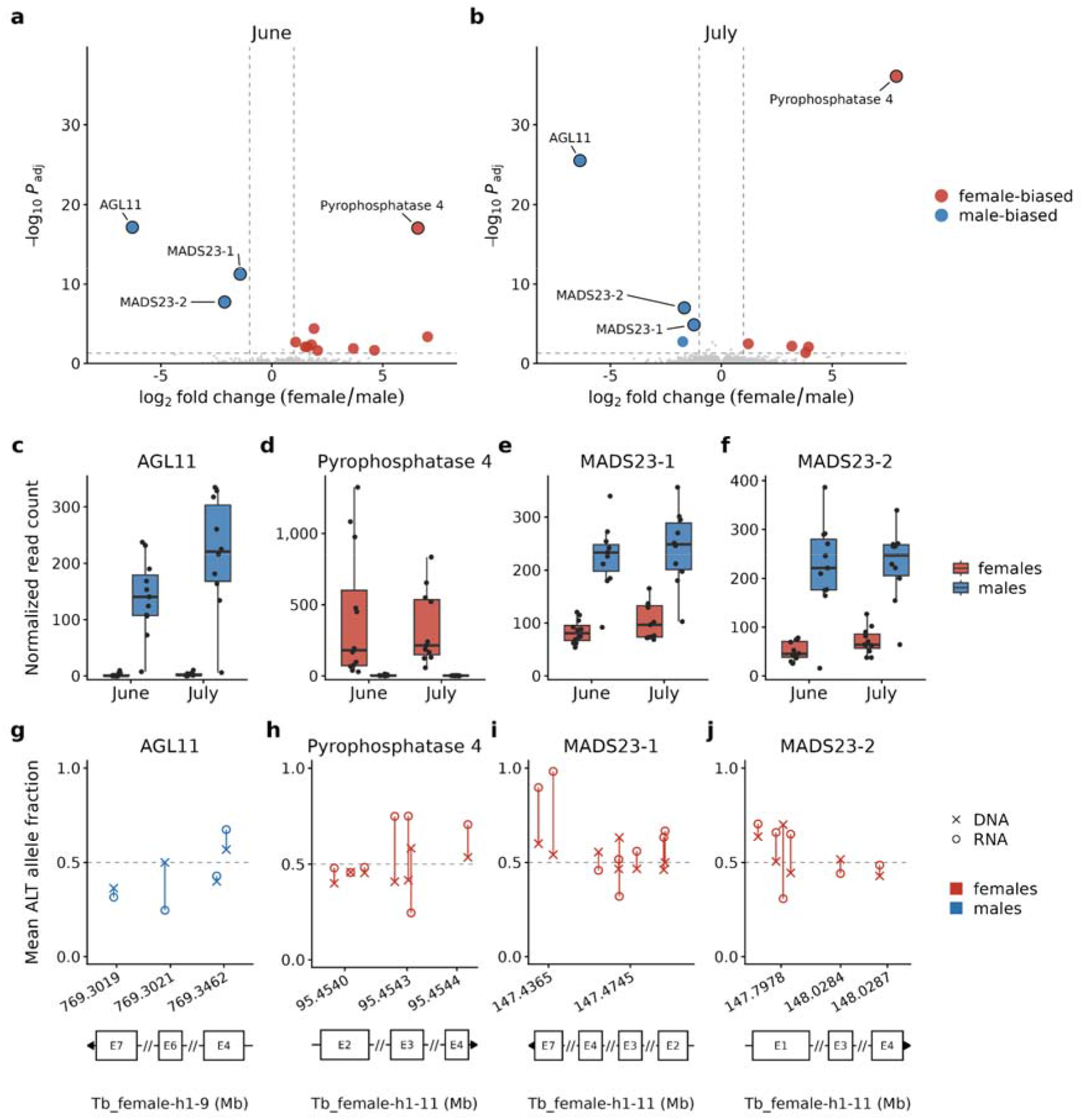
The MADS-box gene AGL11 shows male-specific expression and may act as a downstream factor of sex determination. **a,b** Volcano plots showing 21,153 or 21,235 expressed genes (count > 0 in at least 70% of samples in at least one sex) in reproductive buds from June 2026 (a) and July 2026 (b). Significantly differentially expressed genes in female versus male samples (adjusted PO<O0.05 and absolute log2 fold change > O1.0) are depicted in blue or red. Differential gene expression analysis was performed based on mappings of RNA sequencing reads versus Tb_female-h1. **c-f** Box plots with normalized read counts of female and male samples from June or July 2026 for the genes *AGL11 (g34939)*, Pyrophosphatase *4 (g7677), MADS23-1 (g8053)*, and *MADS23-2 (g8054)*. One male individual exhibited low expression of *AGL11, MADS23-1*, and *MADS23-2*, possibly due to sampling of vegetative buds, which can be difficult to distinguish from reproductive buds at this early stage of development. Boxes indicate the interquartile range with the whiskers extending this range no more than 1.5 times and the center depicting the median. **g-j** Mean fraction in all female or male individuals of the alternative allele (ALT) at selected heterozygous SNP positions (chromosomal positions in exons, marked by E and the exon number) called in short read data from DNA sequencing (DNA) or RNA sequencing (RNA) mapped against Tb_female-h1.

## DISCUSSION

The newly presented haplotype-resolved and chromosome-level genome assemblies for two *T. baccata* individuals appear to be of comparable quality to the most recent genome assembly of *T. wallichiana*^23^. However, our four assemblies representing two female and two male haplotypes provide a unique opportunity to study sex determination, because any X-, Y-, Z- or W-chromosomal sequences should be present. We thus additionally generated short-read resequencing data of 50 female and 50 male individuals to allow extensive reference-free and reference-based genomic analyses to study the genetic basis of sex determination in *T. baccata*.

Sex-specific *k*-mers, which are independent of a reference genome, have proven powerful for identifying the heterogametic sex and for locating SDRs in a range of dioecious plants, from the first identification of a sex-determining factor in persimmon^32^ to the elucidation of the sex-determining system of a lineage that is sister to all other flowering plants, represented by *Amborella trichopoda*^30^. Also, re-analysis of genomic data for the fish *Takifugu rubripes*, in which a single SNP is responsible for sex determination, successfully detected the SDR using *k*-mers^33^. These examples demonstrate that irrespective of the size and genomic context of an SDR, comparisons between relatively small groups of females and males (e.g., 6 vs. 9 for *Amborella* from across its range, or 13 vs. 14 for Fugu) can yield unambiguous results. Our analyses of *Ginkgo biloba*, however, indicate that in species with larger genomes this may not be the case. Unambiguous *k*-mers were only identified when using groups of at least 25 individuals (Fig. 3). The 10 Gb genome appears to result in many common presence-absence variants. In fact, adding even a single *T. baccata* genome assembly to a pangenome graph highlighted 2 billion base pairs of non-core sequence (Supplementary Fig. 9). With this amount of presence-absence variation, clustering by sex is much more likely to occur purely by chance.

Strikingly, using our *Taxus* resequencing data of 100 individuals, we did not identify any sex-specific *k*-mers even though we used different sets of individuals and applied various thresholds (Fig. 3). If sex was nevertheless determined genetically in *T. baccata*, this lack of signal in the reference-free *k*-mer analyses could be due to high genetic differentiation within the group of heterogametic individuals. In larger groups needed due to the large genome size, the causal sex-linked variants might be masked by secondary mutations, while in smaller groups the signal remains non-specific. However, this potential problem should be solved by mapping the short reads to the genome assemblies, which does not rely on exact sequence matches (although it is restricted to genomic regions with proper mapping). Sex-specific sequences should then be identified by sex-specific mapping, a single causal variant in a functional element by a sex-specific signal in a GWAS. Remarkably, neither GWAS nor coverage analyses considering the two female and two male haplotypes identified any sex-linked SNPs or sex-specific sequences (Fig. 4c,d,f,g; Extended Data Fig. 6).

There are only a few additional scenarios, in which an SDR might escape detection. The sex-linked sequence could be part of a repeat array with copy-number variation (CNV) controlling sex. Neither the repeat itself nor its boundaries would be unique to one sex. This could be addressed with a *k*-mer based-GWAS. However, due to the size of the *T. baccata* genome, full *k*-mer-by-sample matrices are not feasible and possible solutions introduce their own limitations^34^. A threshold-based sex-switch controlled by CNV has not been reported for any dioecious plant species so far. Another theoretical scenario may involve a cyto-nuclear interaction, in which a male-(or female-) sterility mutation is inherited through the cytoplasm with an interacting restorer locus in the nuclear genome. However, known examples of cyto-nuclear sex determination involve hermaphrodites, such as gynodioecy in some strawberries^35^. A derived system with stable dioecy would require the evolution of a second nuclear locus. Finally, in the case of polygenic sex determination, in which each locus involved would only show partial sex linkage, our GWAS with 100 samples would not have sufficient statistical power. Again, no example of polygenic sex determination has been reported so far in a dioecious plant species and such a system appears difficult to reconcile with a stable 1:1 sex ratio.

Many well-studied dioecious angiosperms already highlight the remarkable diversity of sex-determining systems and their molecular mechanisms^1,8,36^. In asparagus and kiwifruit, for example, two genes on the Y chromosomes determine sex^37,38^. In persimmon and poplar, single factors on the Y or W chromosomes control sex determination, which in poplar may explain rapid turnover and repeated change of heterogamety^32,39^. Finally, in hops and cannabis, the balance between the X chromosome and the autosomes is responsible for dioecy^3,4^. Since none of those examples would have escaped detection by our analyses, the stage is set for yet another exciting discovery.

In conclusion, our data point to an unconventional basis of sex determination in a dioecious plant species, potentially without sex-specific DNA sequence variation. Future work in *T. baccata* should focus on the analyses of epigenetic marks and additional transcriptional signatures including non-coding transcripts that differentiate females from males. The genome assemblies of both a female and a male *T. baccata* individual presented here provide a solid foundation for such investigations.

## MATERIAL AND METHODS

### Sample collection

*T. baccata* samples used for high molecular weight (HMW) DNA extraction and subsequent PacBio HiFi sequencing or for Hi-C Illumina sequencing were collected in the arboretum of the Thünen Institute of Forest Genetics (Großhansdorf, Germany). Specifically, needles were collected in October 2022 from one male (TXBAC_7_1, Supplementary Table 3) and one female reference individual (TXBAC_17_1, Supplementary Table 3) whose phenotypic sexes were repeatedly assessed over the course of the two previous years. These samples were frozen directly after collection in liquid nitrogen, and stored at U70 °C.

For whole genome (short-read) sequencing, *T. baccata* needle samples were collected or acquired from various sources, including arboreta, botanical gardens and natural reserves (Supplementary Table 3) and stored at −20 °C. Needle samples from Poland were collected with permission of the Regional Directorate for Environmental Protection, Bydgoszcz, Poland (WOP.6400.37.2022, WOP.6205.72.2022.KLD, WOP.6205.73.2022.KLD, WOP.6400.36.2022.MKW).

Tissue samples of the male and female reference individual destined for RNA extraction and subsequent PacBio Iso-Seq were collected in the arboretum of the Thünen Institute of Forest Genetics in Großhansdorf, Germany. Needle samples were collected in May 2023 and frozen in liquid nitrogen directly after collection. Cambium samples were collected in June 2023 near the base of the trunk, using a 16 mm diameter hollow punch to remove a part of the bark and scraping cambium tissue off the removed bark piece with a sharp knife. Root samples were collected in June 2023 by digging up thin roots near the tree trunk. Roots were cut off, briefly washed in tap water and dried with paper towels. Buds were collected in November 2022.

*T. baccata* samples destined for RNA extraction and subsequent short-read RNA sequencing were collected from buds in early stages of development by either plucking them off directly or scraping off the buds from shoots with a knife. Brachyblast buds were targeted as far as the bud identity could be asserted. In June 2023, buds were collected from three male and three female individuals (Supplementary Table 3), including the two reference individuals, in the arboretum of the Thünen Institute of Forest Genetics in Großhansdorf, Germany. The resulting short read data were included in the structural annotation of the four assemblies. Additional bud collections were performed for short-read RNA sequencing in the scope of differential gene expression analysis. For this, buds were collected in the arboretum of the Thünen Institute of Forest Genetics at two time points: (i) on June 17-18, 2026 from 12 female and 11 male individuals as well as (ii) on July 8-9, 2026 from the same individuals with the exception of one female and one male individual (Supplementary Table 3).

Species identity of all samples was confirmed using the species markers TA_ITS, TA_InDel1, TA_InDel2, TA_cox1 (all four markers in ref.^27^) and trnL-F^40^. In the case of root samples, sample identity was further tested via microsatellite analysis to confirm the association between root sample and the respective reference tree. The phenotypic sex of a sample was assigned based on observed strobili and/or arils. When needed, cone buds were assessed under a binocular to avoid confusion with vegetative buds.

### Microsatellite analysis

To ensure genetic distinctiveness among the sampled individuals, all samples destined for whole genome sequencing (WGS) were genotyped with the microsatellite markers Tax36, Tax92 (both in ref.^41^), TS09^42^, Ma-14186-166D^43^, as well as two novel SSR markers named TABAC01 (for: 5’-TATGTGCCTAGGCGTTAGTC-3’, rev: 5’-TTGTAGGTTGATAGACAAATGGA-3’; based on a gSSR (TCC)_7_ on chromosome 2 of *T. chinensis*^44^) and TABAC02 (for: 5’-TTTGGCACTAAACTAAACACATG-3’, rev: 5’-TAATTGTGTTCTCCCTAATAAGG-3’; gSSR (CTT) on chromosome 6 of *T. chinensis*^44^). Genotyping was performed by PCR and subsequent fragment size analysis. Forward primers were tailed at their 5’-end and dye-labeled DNA fragments with complementary sequences to the tails were added to the PCR mix (Supplementary Tables 13 and 14). Forward primers of markers Tax36, TS09, TABAC01 and TABAC02 were tailed with the adapter sequence 5’-CAGGACCAGGCTACCGTG-3’, marker Tax92 with 5’-GCCTTGCCAGCCCGC-3’ and marker Ma-14186-166D with 5’-CGGAGAGCCGAGAGGTG −3’. The PCR reaction setups and PCR programs are given in Supplementary Tables 13 and 14, respectively. PCR products were analyzed on a CEQ 8000 Genetic Analysis System (Beckman Coulter) and evaluated using GeneMarker v3.0.1 (SoftGenetics). When fragment length signals showed stutter bands, the highest peak was chosen. When only one peak was visible, the sample was considered to be homozygous for this marker. Potential clones were identified by pair-wise comparison of fragment sizes (i.e. alleles) using Excel (Microsoft). For heterozygous alleles, the fragments were sorted by size. Each fragment was then compared with the respective fragments of all other samples.

### PacBio HiFi sequencing and Hi-C of the male and the female Taxus baccata reference tree

Nuclei preparation from sample tissue, high molecular weight genomic DNA extraction, Pacific Biosciences (PacBio) HiFi library preparation and sequencing for the samples used in the construction of the reference genomes in this study were previously described^45^ (corresponding to sample IDs 2,6 and 12 in Table 7 of that publication). In short, a total of 2.3 g (over two extractions) and 1.6 g of snap-frozen *T. baccata* needles of the male and female sample, respectively, were used as previously described^45^ to retrieve a total of 26.45 µg (from male sample) and 21.2 µg (female sample) of HMW gDNA. Yield and quality of the extracted gDNA were assessed via NanoDrop and Qubit (BR assay) measurements (Thermo Fisher Scientific) and the fragment lengths were determined on the Femto Pulse (Agilent). HMW gDNA was then sheared to approximately 20 kbp with the MegaRuptor^TM^ (Diagenode). Using 3 µg of high-molecular weight DNA as input, PacBio HiFi libraries were prepared according to the protocol “Preparing whole genome and metagenome libraries using SMRTbell® prep kit 3.0” (Pacific Biosciences) and size selected for fragments larger than 8 kbp with the BluePippin Instrument (SAGE) or larger than 5 kbp with AMPure beads (Beckman Coulter) according to the library preparation protocol. Sequencing was performed on a total of 24 SEQUEL II SMRT™ Cells with 30-hour movies (10 for the female sample, 14 for the male sample), using the Sequel II Binding Kits 3.2 and the Sequel II sequencing kit 2.0 (Pacific Biosciences). Raw reads were processed using ccs v6.4.0 (https://github.com/PacificBiosciences/ccs), actc v0.6.0 (https://github.com/PacificBiosciences/actc) and deepconsensus v1.2.0^46^. Sequencing resulted in 410.02 Gb and 359.32 Gb circular consensus sequence reads for the male and female individual, respectively.

For Hi-C data, nuclei from sample tissue were extracted as described in ref.^45^. Hi-C libraries were prepared for each sample using the Arima-HiC Kit (Arima Genomics, Material Nr. A510008) and KAPA HyperPlus Kit (Kapa Biosystems) according to the manufacturer’s protocols. For the female sample, a 0.78X AMPure bead (Beckman Coulter) size selection was performed to eliminate < 200 bp fragments. Both libraries have been sequenced to ∼63× (Tb_female) or ∼50× (Tb_male), with 200 cycles on an Illumina NovaSeq 6000 platform.

### Whole genome sequencing of 103 Taxus samples

DNA extraction for WGS was performed as previously described^47,48^ (the latter protocol for difficult samples; see Supplementary Table 3). DNA was checked via NanoDrop and Qubit (BR Assay) measurements (both: Thermo Fisher Scientific). DNA was sheared to 400 bp fragments (on average) using a Covaris LE220 ultrasonicator (Covaris), and 250 ng of sheared DNA was used as input for the KAPA HyperPlus Kit (Kapa Biosystems) to construct paired-end Illumina libraries, using 5 amplification cycles. Sequencing to approximately 20× coverage as 2×150 bp paired-end reads was performed with NovaSeq S4 flowcells and v1.5 reagents on an Illumina NovaSeq 6000 platform (Illumina).

### RNA extraction and sequencing

Frozen sample material was ground using mortar and pestle and RNA was extracted following the Spectrum™ Plant Total RNA-Kit (Sigma-Aldrich) protocol B instructions (including the optional DNase digestion step) with the addition of 30 mg Polyclar (SERVA) to the lysis buffer for each sample. RNA quality was assessed via NanoDrop, Qubit (BR assay) measurements (both: Thermo Fisher Scientific) and Bioanalyzer, using a Plant RNA Nano assay (Agilent).

For PacBio Iso-Seq sequencing, Iso-Seq libraries of total RNA of different tissues of the male and female reference sample were generated according to the PacBio protocol ‘Preparing Iso-Seq® libraries using SMRTbell® prep kit 3.0 (PN 102-396-000 REV02 APR2022)’. In brief, 300 ng total RNA was reverse-transcribed with the NEBNext® Single Cell / Low input cDNA synthesis and amplification module, with barcoded cDNA primers in the cDNA amplification step. Barcoded cDNAs from needle, bud, and root samples were pooled separately for each of the two reference individuals. For cambium samples, the barcoded cDNAs from the two individuals were pooled together. PacBio sequencing adapters were then ligated to each pool, generating three libraries that were sequenced on three PacBio Sequel II SMRT cells for 30 hours with the Sequel II Binding Kits 3.2 and the Sequel II sequencing kit 2.0. After circular consensus calling of the raw sequencing reads, high-quality isoforms were called and clustered with the default SMRTlink Iso-Seq pipeline (v11 and v13).

For short-read RNA sequencing in June 2023 (buds from three male and three female individuals; see “Sample collection”), libraries with an average insert size of 335 bp were constructed using the NEBNext® Ultra™ II Directional RNA Library Prep Kit for Illumina® (poly-dT enrichment workflow) (New England Biolabs). Sequencing to approximately 60 million reads as 2×150 bp paired-end reads was performed with NovaSeq S4 flowcells and v1.5 reagents on an Illumina NovaSeq 6000 platform (Illumina).

For short-read RNA sequencing in June 2026 (12 females, 11 males) and July 2026 (11 females, 10 males; see “Sample collection”), library preparation and sequencing were performed by Novogene. mRNA was enriched from total RNA by Poly-A-selection and strand specific, dual indexed cDNA libraries were prepared by using the dUTP method. Short-read RNA sequencing was done with Illumina NovaSeq X Plus resulting in approx. 34 million paired reads (2×150 bp) per sample.

### Genome assembly

The assembly procedures of the female and male individuals differed slightly: For the reference assembly of the female sample, assemblies were generated with Hifiasm^49,50^ v0.18.9-r527. Remaining haplotype duplications were manually removed, guided by purge_dups^51^ v1.2.6 (https://github.com/dfguan/purge_dups), to avoid over-purging in repetitive regions. Scaffolding of the contigs was performed by YaHS^52^ v1.2a.1. Chloroplast and mitochondrial contigs were identified with Tiara^53^ v1.0.3 and MitoHifi^54^ v3.2.1, respectively, and manually removed. BlobToolKit^55^ v4.4.5 was used to identify and remove contaminations. The male reference assemblies were generated with Hifiasm^49,50^ v0.19.8-r603, duplications were removed with purge_dups^51^ v1.2.5 (https://github.com/dfguan/purge_dups) and scaffolded with YaHS^52^ v1.2.

All scaffolds were then manually curated in PretextView v1.0.3 (https://github.com/sanger-tol/PretextView), GRIT_Rapid (https://gitlab.com/wtsi-grit/rapid-curation, commit 1a3d79a8), HiGlass^56^ v0.10.4, and the DAmar pipeline (https://github.com/MartinPippel/DAmar; branch: master-v1, commit 1e428cddaa79fee04edbcbc41bcca1a12a014b11).

Assembly completeness was assessed with BUSCO^57^ v5.8.2 with the embryophyta_odb12 lineage dataset.

### Generation of a pangenome graph

A pangenome graph with four haplotypes of *T. baccata* was generated using Minigraph-Cactus pipeline^58^. The pipeline was implemented within the Cactus v3.1.2 and was run using the cactus-pangenome command with default parameters. To assess and visualize the growth of the pangenome we used panacus^59^ v0.3.3.

### Structural genome annotation with BRAKER3

To generate structural annotations of the female and male reference assemblies (two haplotypes each), the TETools container v1.90 (https://github.com/Dfam-consortium/TETools) and BRAKER^60–64^ container v3.0.7.6 (https://github.com/Gaius-Augustus/BRAKER) were utilized. First, a library of transposable elements was generated *de novo* by RepeatModeler^65^ v2.0.6 and contained software, including RepeatScout^66^ v1.0.7, Tandem Repeats Finder^67^ v4.09, RECON^68^ v1.08, LTR_retriever^69^ v2.9.0, NINJA^70^ v1.00-cluster_only, CD-HIT^71^ v4.8.1, MAFFT^72^ v7.471, HMMER v3.4 (http://hmmer.org/), GenomeTools^73^ v1.6.4, and a selection of UCSC utilities (see https://github.com/Dfam-consortium/TETools). For this step, the LTR Structural Analysis option (“-LTRStruct”) was enabled, apart from this default options were used. Based on this library, the reference assemblies were soft-masked with RepeatMasker v4.1.7-p1 (http://repeatmasker.org) using default parameters (except for the soft-mask setting “-xsmall”). Subsequently, the assemblies were soft-masked further with Tandem Repeats Finder v4.09^67^, using the parameters “2 7 7 80 10 50 500 -d -m -h”.

Iso-Seq transcript data for usage in BRAKER was prepared separately to account for large introns in the target species. To estimate the maximum intron size of the *Taxus* genome while avoiding false positives (i.e., wrong alignments that erroneously indicate very large introns), Iso-Seq transcripts were mapped to the female reference assembly haplotype 1 using minimap2^74,75^ v2.28 multiple times with the parameters “-uf -a -x splice:hq” and varying “-G” parameter settings. Mappings were then filtered to exclude alignments with the “secondary” or “supplemental” flag, and alignments with a mapping quality < 40. From each mapping, the longest N-operation in the CIGAR string (indicating a skip in the reference compared to the read, i.e., possibly an intron) was determined and plotted against the used -G parameters. The resulting plot showed a region in which the maximum N-operation length stagnated even under increasing “-G” parameter settings while at even higher “-G”, maximum N-operation length rose quickly again (i.e., the plot formed a plateau). The “-G” parameter which corresponded to the middle of this region was then used for the mappings in BRAKER.

The soft-masked reference assemblies were each used as genome data input for two BRAKER^61–64^ workflows: (i) BRAKER3 with short-read RNA input and (ii) BRAKER3 with long-read RNA input^60^. For (i), RNA data input consisted of the short-read RNA sequencing data generated for this study based on the bud sample collection in June 2023 (see sample collection), and a public dataset of a *T. baccata* transcriptome^76^ (SRA accessions SRX2999991, SRX2999990). For this workflow, v3.0.7.6 of BRAKER was used.

For (ii), PacBio Iso-Seq transcripts that were generated for this study were mapped to the respective reference genome with minimap2^75^ v2.28 using the parameters “-G 500k -uf -a -x splice:hq”. The resulting mapping dataset was used as RNA input. In both workflows, protein input data consisted of the publicly available OrthoDB (https://www.orthodb.org/) partition Viridiplantae_v12^57^ and a UniProt^77^ (https://www.uniprot.org/) dataset of 10991 peptide sequences (using the filters “(taxonomy_id:25628) AND (existence:3)”, downloaded on 2024-11-27). For this version of the workflow, the modified BRAKER docker container for Iso-Seq input (docker://teambraker/braker3:isoseq) was used. In addition to BRAKER itself, the following software was used in the pipeline: AUGUSTUS^78,79^ v3.5.0, Spaln^80^ v2.3.3f, StringTie2^81^ v2.2.1, GFF utilities^82^ v0.12.7, TSEBRA^83^ v1.1.2.5, SAMtools^84^ v1.13, BamTools^85^ v2.5.1 and BEDTools^86^ v2.30.0. Merging of the output of the two workflows was performed using TSEBRA^83^ v1.1.2.5 with the “--filter_single_exon_genes” option. The two workflows each used new separate species profiles for AUGUSTUS.

Descriptive statistics of the predicted gene sets were generated with a custom awk script (see Code Availability). BUSCO scores of the gene sets were generated via BUSCO^57^ v5.8.2 in conjunction with the embryophyta_odb12 lineage dataset. Gene predictions were visually inspected in CLC Genomics Workbench v24.0.2 (QIAGEN) by importing the reference genome FASTA file, annotation GTF and the intermediate “hintsfile” GTF (containing extrinsic evidence from RNA and protein data, i.e., intron, CDS, start and stop codon hints) files, and RNA mapping BAM files, converting these files into “tracks” and overlaying these datasets as “track list” in CLC.

To assess annotation consistency across haplotypes and identify unique and shared genes, we used Liftoff^87^ v1.6.3. Liftoff transfers annotations by aligning gene sequences from the annotated query genome to the target genome. We applied sequence identity and alignment coverage thresholds of 0.5. The resulting gene-presence sets were visualized using UpSetR^88^ v1.4.0.

### Alternative structural genome annotation with Helixer

Annotations supplementary to those generated by BRAKER3 as described above were predicted *de novo* using Helixer^89^ v0.3.6. Unmasked assemblies were supplied as inputs and the --lineage parameter was set to “land_plant”. Coding sequences and amino acid sequences were extracted by parsing the Helixer-generated GFF3s and the corresponding genome assemblies using gffread^82^ v0.12.7.

### Functional genome annotation

Protein-coding sequences were further annotated using the EnTAP^90^ container v2.2.0. EnTAP was used in conjunction with the frame selection step with TransDecoder v5.7.1 (https://github.com/TransDecoder/TransDecoder/), the similarity searching step with DIAMOND^91^ v2.1.8 and the NCBI RefSeq databases (https://www.ncbi.nlm.nih.gov/refseq/) for plant, fungi and bacteria, and the ontology analysis step with EggNOG^92^, using the eggnog-mapper^93^ v2.1.12.

### Gene Ontology overrepresentation analyses in sets of haplotype-specific genes

For each haplotype we performed an overrepresentation analysis (ORA) with the R package ClusterProfiler^94^ v4.10.6 in R v4.5.0. The test set comprised all haplotype-specific genes. Background genes were defined by all genes of the same haplotype with at least one Gene Ontology annotation. Terms of all three aspects of the Gene Ontology (biological process, molecular function and cellular component) were collectively tested. Only terms that were assigned 10 to 500 background genes were considered for testing. The enrichment was tested with a hypergeometric test and *p*-values were corrected for multiple testing using the Benjamini-Hochberg method^95^. Terms with an adjusted *p*-value < 0.05 were considered significantly overrepresented.

### Identification of structural variants in the four haplotype assemblies

To compare the haplotype assemblies and identify large structural variants, alignments between the assembly sequences were generated with mm2plus^96^ v1.1 using the parameters “-x asm5 -k 28 -I 50G -c --eqx“. The mappings were then processed with SyRI^97^ v1.7.0 and visualized with Plotsr^98^ v1.1.1. A complementary approach using protein orthologs (generated by structural genome annotation, see above) was conducted with GENESPACE^99^ v1.3.1.

### Validation of selected structural variants in Hi-C contact maps

We cross-validated the following structural variants by inspecting the Hi-C contact maps: an ∼97 Mb inversion (Tb_female-h1-8: 284.2–381.3 Mb) and an ∼13 Mb inversion (Tb_female-h1-8: 381.3–394.6 Mb) adjacent to each other on chromosome 8 and an ∼46 Mb inversion on chromosome 3 (Tb_female-h1-3: 420.5–466.4 Mb). For this, Hi-C reads of Tb_female or Tb_male, respectively were mapped to each female or each male haplotype assembly, so that reads from both homologous chromosomes are shown on the same reference. Next, we generated cooler files from the Hi-C mappings using cooler v0.10.4 (binning contacts at 10 kb, aggregated to multiple resolutions) and visualized the interactions using ggplot v4.0.4 as detailed in Extended Data Figs. 4 and 5.

### Quantification of annotation features affected by genomic variants

Genomic intervals of structural annotation features (genes, introns and exons) and imputed complementary regions (promoter regions as up to 5 kb upstream of each gene interval, and intergenic regions as all bases that are not part of any other feature), were prepared based on the structural annotation with BRAKER3 in BED format using bedtools^86^ v2.31.1. Similarly, the intervals of genomic variation features based on SyRI output were prepared, so that for each haplotype, the genomic variants compared to the three other haplotypes were written in BED according to their size in three categories: SNPs, small SVs (<= 50 bp) and large SVs (>50 bp). For each combination of genomic feature type and genomic variant type, the number of genomic features with at least one bp overlap with the genomic variant were determined with bedtools^86^ intersect v2.31.1. The fractions of affected features were then calculated in R^100^ v4.5.0 and visualized with the ggplot2^101^ package v3.5.1.

### K-mer analysis

To identify sex-specific sequence regions, a binary presence–absence *k*-mer analysis^30^ was applied using the trimmed WGS reads and BiTUGA^34^ with a *k*-mer size of 31. The canonical *k*-mer pools for both groups (50 female and 50 male samples) were generated and then filtered under multiple criteria using the --group-min and --group-max options in BiTUGA pass II, with other filters disabled.

For the second approach, 10 sets of five female and five male samples were generated from the original sample set with no duplicates. Additionally, six sets of individuals were generated in which each set contained all samples from one autochthonous population. The samples from Jelenia Góra and Wierzchlas were considered as one autochthonous population in this analysis due to low geographical distance of about 10 km. For these sets, new *k*-mer databases were generated like above, and filtering was performed so that one individual from each group may diverge with regard to the prevalence filters. From the BiTUGA output, we then considered both the “post-bias” *k*-mer counts as well as the “perfect contrast” counts (that correspond to the absolute filtering without divergence). Determination of consistency was performed based on the BiTUGA output files using a custom python script (see code repository).

For *k*-mers from *Ginkgo biloba*, we first compared sample IDs from NCBI BioProject PRJNA478810 with those listed in Supplementary Table 4 from Zhang et al.^102^. We only included samples sequenced on the BGISEQ-500 platform, with an average spot length of 50, a sequencing depth between 5 and 9, and a mapping rate of at least 90%. With 221 samples remaining, we randomly sampled 50 female samples and 50 male samples to generate our final sample set and downloaded the respective WGS reads using the sra toolkit v3.0.7 (https://github.com/ncbi/sra-tools). The above analyses were then performed with the *Ginkgo* read data. Read trimming was not performed for these reads due to low adapter occurrence as determined by FastQC v0.12.1 (https://www.bioinformatics.babraham.ac.uk/projects/fastqc/; with BGI adapter sequences taken from https://github.com/OpenGene/fastp/blob/master/src/knownadapters.h).

Visualization of all analysis results was performed with the ggplot2^101^ package v3.5.1 in R^100^ v4.5.0.

### Mapping of WGS data

WGS paired-end reads were trimmed with Trimmomatic^103^ v0.39 using the settings “ILLUMINACLIP:$adapter:2:30:10:1:true”, “SLIDINGWINDOW:4:15” and “MINLEN:60”. The adapter sequence file ($adapter) corresponds to the TruSeq3-PE-2.fa file in the trimmomatic github repository (https://github.com/usadellab/Trimmomatic/blob/main/adapters/TruSeq3-PE-2.fa). Before and after trimming, the reads were assessed using FastQC v0.12.1 and MultiQC^104^ v1.27.1. Unpaired trimmed reads were excluded from further analysis. The remaining trimmed reads were mapped onto reference genome assemblies (four haplotypes in total) using bwa-mem2^105^ v2.2.1. Mapping quality was assessed with Qualimap^106^ v2.3. Read mappings corresponding to the same biological sample were then merged using the SAMtools^84^ v1.12 function “merge” with the options “-r -t SQ”. Potential duplication artifacts (i.e., reads originating from the same DNA fragment) in the merged files were marked with MarkDuplicates v3.1.0 of the Picard tool suite, using the option “--OPTICAL_DUPLICATE_PIXEL_DISTANCE 2500”.

### Variant calling

Variant calling was performed on the read mappings using bcftools^84^ mpileup and call with the options “--ploidy 2 -m -v”. From the resulting VCF files, indels were excluded from all further analyses.

### Population structure analysis

For PCAs (intended to correct for population structure during GWAS), subsets of SNPs were generated, containing 140853, 101647, 128570 and 134459 SNPs in the sets of Tb_female-h1, Tb_female-h2, Tb_male-h1 and Tb_male-h2, respectively. For these sets, descriptive statistics were generated and visualized using bcftools^84^ v1.21 and the ggplot2^101^ package v3.5.1 in R^100^ v4.4.2.

Based on these visualizations and the GATK recommendations for hard-filtering of germline variants (https://gatk.broadinstitute.org/hc/en-us/articles/360035890471-Hard-filtering-germline-short-variants), the bcftools-generated VCF files were filtered to exclude variant sites as follows: (1) ‘INFO/FS < 0.000001 || INFO/MQ < 40 || INFO/MQBZ < −12.5 || RPBZ < −4 || COUNT(GT=”mis”) > 0.1 * N_SAMPLES’, (2) Quality-by-depth metric less than 5 (these were calculated according to the documentation for GATK’s QualByDepth), (3) non-biallelic SNPs. Filtered VCF files were then imported into PLINK^107^ v1.9b7 and further filtered to exclude variants that deviated from Hardy-Weinberg equilibrium at p < 1e-8. Fully filtered variant sets were then subjected to variant pruning by linkage disequilibrium (LD). For this, pairwise LD statistics were generated in PLINK v1.9b7^107^ and visualized with the ggplot2^101^ package v3.5.1 in R^100^ v4.4.2. Based on an LD decay plot, a cutoff was determined and variants were pruned to exclude variants with a correlation of r² > 0.2 in 100kbp windows (step size 20) with PLINK^107^ v1.9b7.

### GWAS

For GWAS, subsets of SNPs were generated from the unfiltered VCF file (without InDels) with bcftools^84^ v1.21 to exclude (1) non-biallelic SNPs, (2) SNPs with a minor allele frequency of less than 10%, (3) SNPs with a median allelic depth smaller than 4 or larger than 64, (4) SNPs that are missing in more than 10% of samples, after excluding the three problematic samples, i.e., the two *T.* x *media* hybrids and the one *T. baccata* tree with a cosexual branch. A GWAS was performed with GEMMA^108^ v0.98.5 using a linear mixed model (-lmm 2), using phenotypic sex as a binary variable and the first 6 axes of the previously generated PCA as covariates (option-c). All other filtering steps applied by default in GEMMA were deactivated. To visualize GWAS results as Manhattan plots, the ggplot2^101^ v4.0.0 package in R^100^ v4.5.0 was used.

### Analysis of genome-wide F_ST_ and nucleotide diversity

To determine genomic regions with anomalous *F*_*ST*_ or nucleotide diversity (π) that could be indicative of an SDR, we employed clam^109^ v1.1.4. Based on the WGS read mapping depths of the 100 samples, loci with sufficient coverage to ascertain the genotype were determined (minimum 3, maximum 250), and based on the same SNP data used in the GWAS approach, *F*_*ST*_ and nucleotide diversity (π) were calculated for non-overlapping 100 bp windows across all four assemblies, with samples being grouped according to phenotypic sex instead of populations. For plotting, the 100 bp values were combined into non-overlapping 100 kb windows: *F_ST_* as the mean of the 100 bp values, and π per sex as the total number of pairwise differences divided by the total number of pairwise comparisons. π is shown per 100 kb window together with a centered 5 Mb rolling mean. *F*_*ST*_ is shown as the mean and the range of the 100 kb values within each plotting bin (∼2.4 Mb at the printed figure width).

### Analysis of differential mapping coverage

The read coverage across all four haplotypes was determined with mosdepth^110^ v0.3.11, using 100 bp non-overlapping windows and a minimum mapping quality of 20. For each sample, a median coverage of these windows was calculated, and each window’s depth was divided by this median, so that a value of 1 corresponds to normal diploid coverage.

As a quality filter, a window was kept only if at least 25 of the 50 individuals of the sex that must carry every sequence of the assembly, i.e. females for the female assembly including any potential W-specific sequence and males for the male assembly including any potential Y-specific sequence, had a normalized depth of ≥ 0.2. This kept 93–94% of windows per haplotype.

The mean normalized depth of each sex was calculated per window. For visualization, blocks of 1,000 windows were analyzed corresponding to ca. 100 kb. For each block, a chromosome quotient was computed as the summed mean depth of the non-carrying sex (ZZ male or XX female) divided by that of the carrying sex, i.e. male/female on the female assembly, and female/male on the male assembly. The expected quotient is 1 for autosomal sequence, 0 for sequence specific to the W or Y chromosomes, and 2 for Z- or X-specific sequence on the female and male assemblies, respectively.

In addition, every 100 bp window was screened in individual samples for the patterns expected for a sex-specific region, with a normalized depth of < 0.05 counted as absent, 0.25–0.75 as one copy and 0.75–1.25 as two copies. First, we tested for presence of one copy in the carrying sex and absence in the other, to identify potential W- or Y-specific 100 bp windows. Second, we tested for presence of one copy in the carrying sex and two in the other, to identify potential Z- or X-specific 100 bp windows. We used the smaller of the fractions of females and males matching their expected class, so that a truly sex-specific region scores 100%. No single window came close to the pattern expected for a sex-specific region (best scores 58–68%).

The chromosome quotient was plotted along the genome for each haplotype. The best-scoring single window of each screen is marked by a triangle at its own quotient.

### Differential gene expression analysis

Read mapping and quantification of read counts were done with the STAR aligner^111^ v2.7.11a against all scaffolds of the Tb_female-h1, Tb_female-h2, Tb_male-h1, and Tb_male-h2 assemblies with two-pass mode and maximal intron length set to 600000. Differential gene expression analysis was done in R v4.5.0 with the Bioconductor package DESeq2^112^ v1.48.2 with the significance cutoff value alpha set to 0.05. Genes were only tested when 70% of all individuals in one sex had counts > 0. The model only included sex as factor (∼ sex, reference male). Genes were labelled differentially expressed if padj < 0.05 and |log2FC| > 1.

### Candidate gene screening for heterozygous DNA sequence variants and monoallelic RNA expression

Allele-specific expression was analyzed in the genes *AGL11*, *MADS23*-1, *MADS23*-2 and Pyrophosphatase 4 to identify potential monoallelic expression.

DNA and RNA sequencing data were available for 5 female and 5 male individuals. DNA short reads were mapped against all scaffolds of the Tb_female-h1 assembly with bwa-mem2^105^ v2.2.1 and duplicate reads were marked with Picard MarkDuplicates (https://broadinstitute.github.io/picard/) v3.1.0. Variants were called within the genes plus 10 kbp flanking sequences with bcftools v1.22 (mpileup with mapping quality >= 20, base quality >= 20 and maximum depth 1000; call; norm).

Variants with QUAL < 20 were removed and individual genotypes supported by fewer than 5 reads were set to missing. SNPs indicating collapsed paralogous regions were excluded, i.e. SNPs meeting all three conditions: at least 80% of individuals were genotyped, at least 90% of the genotyped individuals were heterozygous and the total read depth across individuals was at least 1.5 times the median depth of all filtered variants. STAR alignments were reduced to gene loci plus 1 kbp of flanking sequence with SAMtools^84^ v1.22.1 and duplicate reads were marked with Picard MarkDuplicates v3.1.0. Alignments of every individual from June and July were merged. Reads crossing intron borders were split with GATK^113^ SplitNCigarReads v4.4.0.0.

To identify monoallelic expression, heterozygous SNPs were selected in the DNA sequencing data of every individual based on the following criteria: GQ >= 20, read depth >= 8, minimum two reads per allele, allele balance between 0.3 and 0.7. Only biallelic SNPs in exonic regions were considered. At the selected SNP positions RNA short reads were counted in the read mappings for each allele with bcftools mpileup, without genotyping. The following filter criteria were used: mapping quality >= 20, base quality >= 20, BAQ deactivated, maximum read depth 10000. PCR duplicates were not counted. Indels were disregarded. For every individual and every SNP, the allele fraction was calculated as follows: AF = nALT / (nREF + nALT). Reads with a third allele were disregarded. For the comparison between DNA and RNA, a SNP was required to have at least 10 reads (REF + ALT) in both DNA and RNA in the same individual.

## ACKNOWLEDGMENTS

This study was funded by the Deutsche Forschungsgemeinschaft (DFG, German Research Foundation) in the scope of the TaxGen project (DFG project 497528752 awarded to B.K.; DFG grant no. KE 916/10-1) and supported by the DFG Research Infrastructure NGS_CC (DcGC, project 407482635) as part of the Next Generation Sequencing Competence Network (DFG project 423957469). NGS library preparation, data production and analyses were carried out at the DcGC Dresden-concept Genome Center - a core facility of the CMCB and Technology Platform of the TUD Dresden University of Technology. D.L. was supported by the DFG project 545520056 awarded to N.A.M. V.R. was supported by the DFG project 544948635 awarded to N.A.M. Additional funding was received by the Institute of Dendrology, Polish Academy of Sciences. We thank K. Groppe, V. Kuhlenkamp, A. Schellhorn, A. Eikhof, I. Burau and gardeners from the Thünen Institute of Forest Genetics for technical assistance, and M. Fladung for helpful comments and discussions. We thank the following contributors from the MPI-CBG and the DcGC for wet lab work: N. Gscheidel (for gDNA preparation and PacBio sequencing), D. Pache (for PacBio sequencing), W. Tan (for Hi-C). We are grateful to M. Krause and J. Gusson Roscito from DcGC for coordination of Illumina sequencing. We thank S. Rust, J. Schöttler (both: Loki Schmidt Botanical Garden Hamburg, Germany) and S. Petersen (Botanical Garden Kiel, Germany) for providing access to the related botanical gardens, and M. Seho (Bavarian Office for Forest Genetics, Teisendorf, Germany) for providing additional samples. We thank T. de Jong (Leiden University, Netherlands) for helpful discussions.

## DATA AVAILABILITY

All read data, assemblies and annotations have been deposited at ENA/GenBank/DDBJ; with Illumina short-read data, Hi-C reads, PacBio HiFi long reads, genome assemblies and annotations accessible under the BioProject accession PRJEB103025; and PacBio Iso-Seq reads and transcriptome assemblies deposited under the BioProject accession PRJNA1554957.

## CODE AVAILABILITY

All scripts (and related source data files) used to assemble the genomes, analyze the data and visualize the results are deposited in https://github.com/daniel-bross/Taxus_genome

## AUTHOR CONTRIBUTIONS

D.B., J.M., N.A.M. and B.K. conceptualized the project.

S.W., N.A.M. and B.K. supervised the project and conceived the methodology.

D.B., E.P.K., H.S. and B.K. collected samples.

S.W. coordinated the sequencing steps.

M.P. and L.U. performed the genome assemblies, M.P. performed the manual curation of assemblies and D.B. and V.R. annotated the assemblies. D.L. performed the comparative annotation and pangenomic analyses.

D.B., D.L., M.M., J.M., M.P., S.K., L.U., N.A.M. and B.K. analyzed the sequencing data.

S.K. and B.K. supervised *k*-mer analyses.

H.S. advised SSR-marker analyses.

D.B., D.L., M.M., M.P., N.A.M. and B.K. conducted the visualization.

D.B., N.A.M. and B.K. wrote the manuscript, and all authors read, edited and approved the manuscript.

E.P.K., S.W., N.A.M. and B.K. acquired the funding.

## EXTENDED DATA FIGURES

**Extended Data Figure 1.**
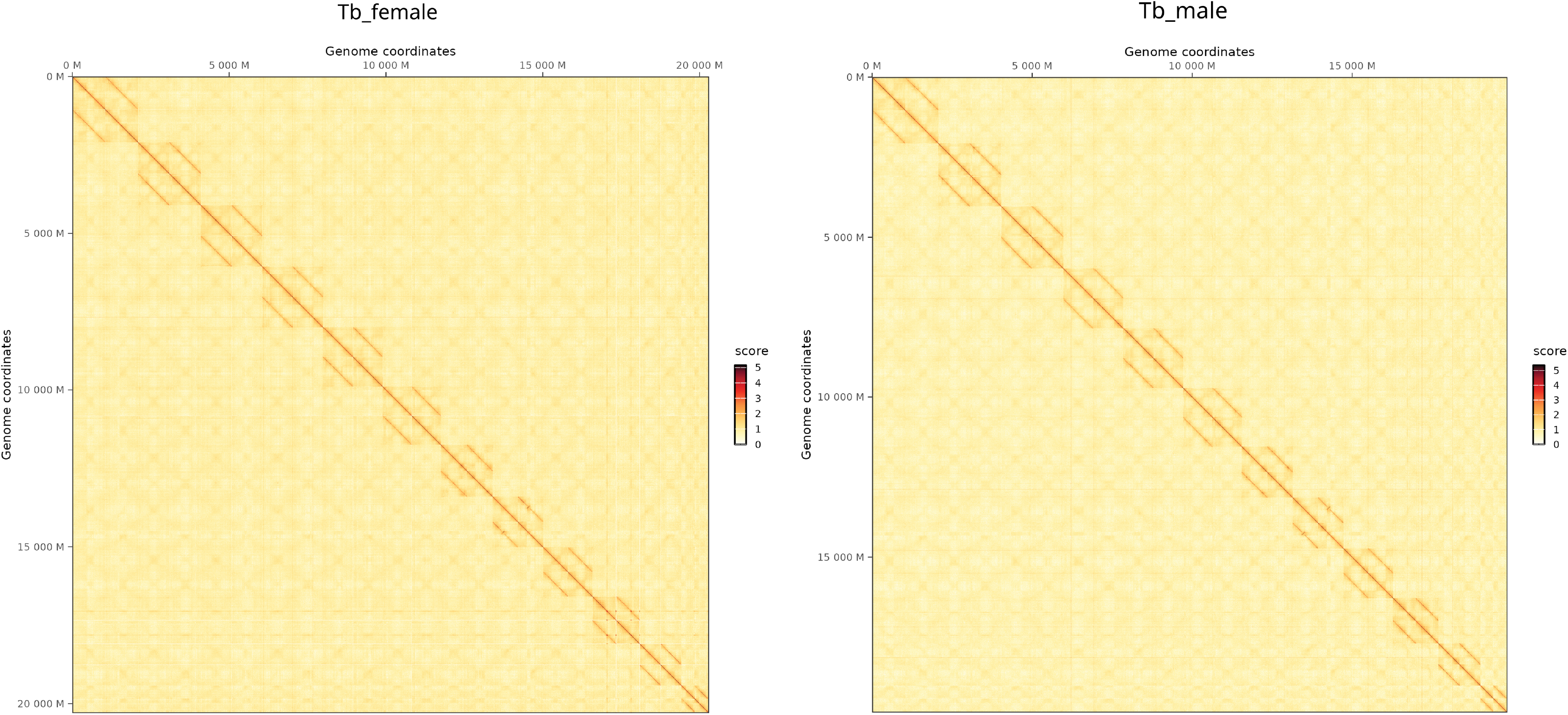
Diploid Hi-C interaction heatmaps of the two assembled haplotypes of each of the sequenced individuals show successful scaffolding into chromosomes. For each individual (female: Tb_female-h1 and Tb_female-h2; male: Tb_male-h1 and Tb_male-h2), the Hi-C reads were mapped to the combined assembly of both haplotypes, retaining reads that map equally well to both haplotypes (MAPQ ≥ 0). Assembled chromosomes 1 to 12 are shown in the order of size, haplotype 1 followed by haplotype 2, on both axes. The blocks along the main diagonal represent the intra-chromosomal contacts of each chromosome; the absence of signal between different chromosomes confirms the successful scaffolding. The inter-haplotype blocks display a strong, distinct diagonal of elevated signal, which mainly reflects read cross-mapping between homologous chromosomes and, to a lesser extent, trans-homolog interactions. Color intensity indicates Hi-C contact frequency. Hi-C contacts (displayed as balanced counts) are shown at a bin size of 2 Mb. The plots were generated with cooler and HiContacts.

**Extended Data Figure 2.**
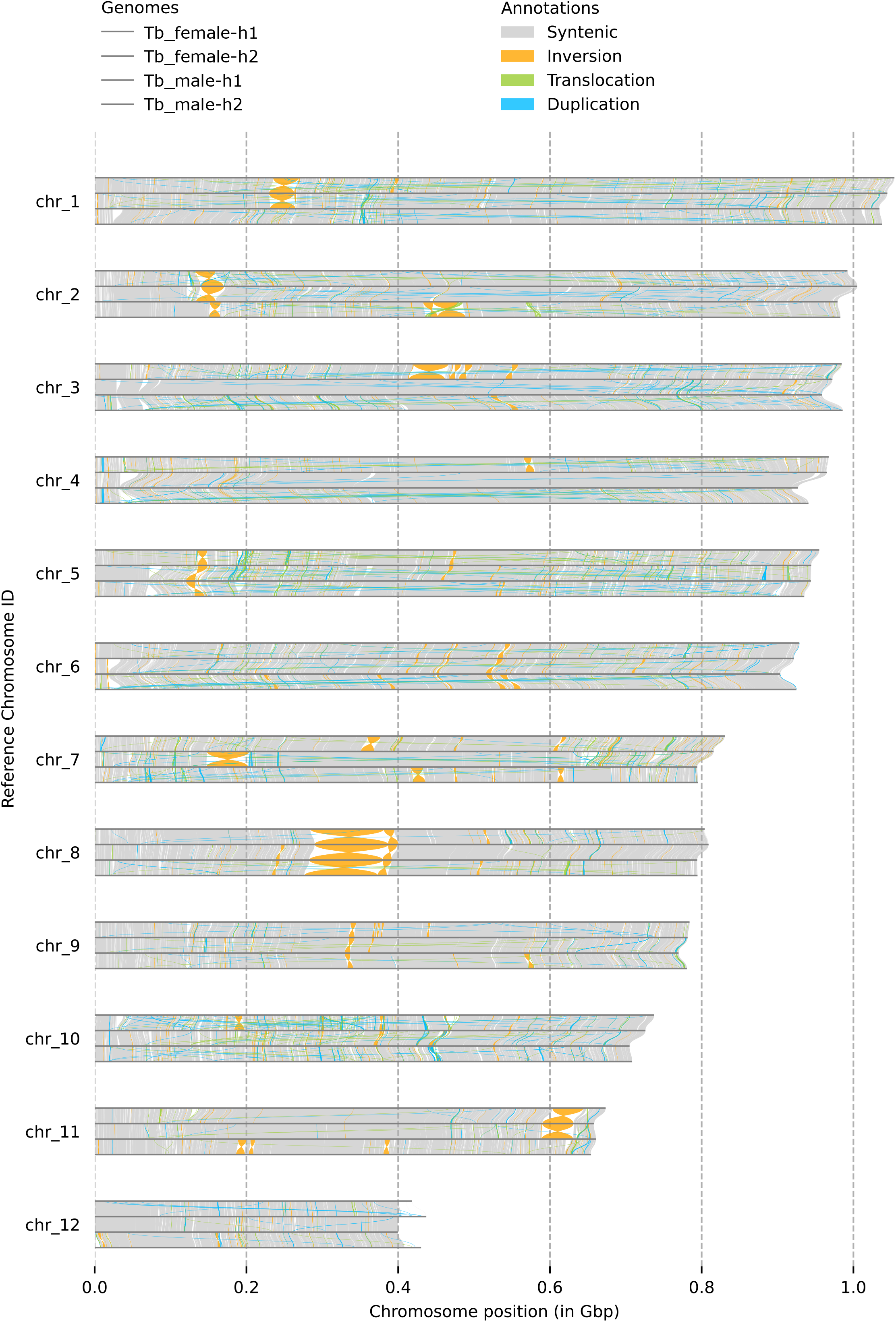
Pairwise sequence alignments and synteny analysis with SyRI highlight large structural variants between the four haplotype-resolved assemblies. For each of the 12 *T. baccata* chromosomes, the assemblies of the four haplotypes are arranged one below the other in the following order: Tb_female-h1, Tb_female-h2, Tb_male-h1, Tb_male-h2. The following structural variants between two adjacent haplotypes are depicted in color code: inversions (orange), translocations (green), and duplications (blue). Syntenic regions between two haplotypes are displayed as grey outlines. Only variants with a size of 10 kb or greater are displayed.

**Extended Data Figure 3.**
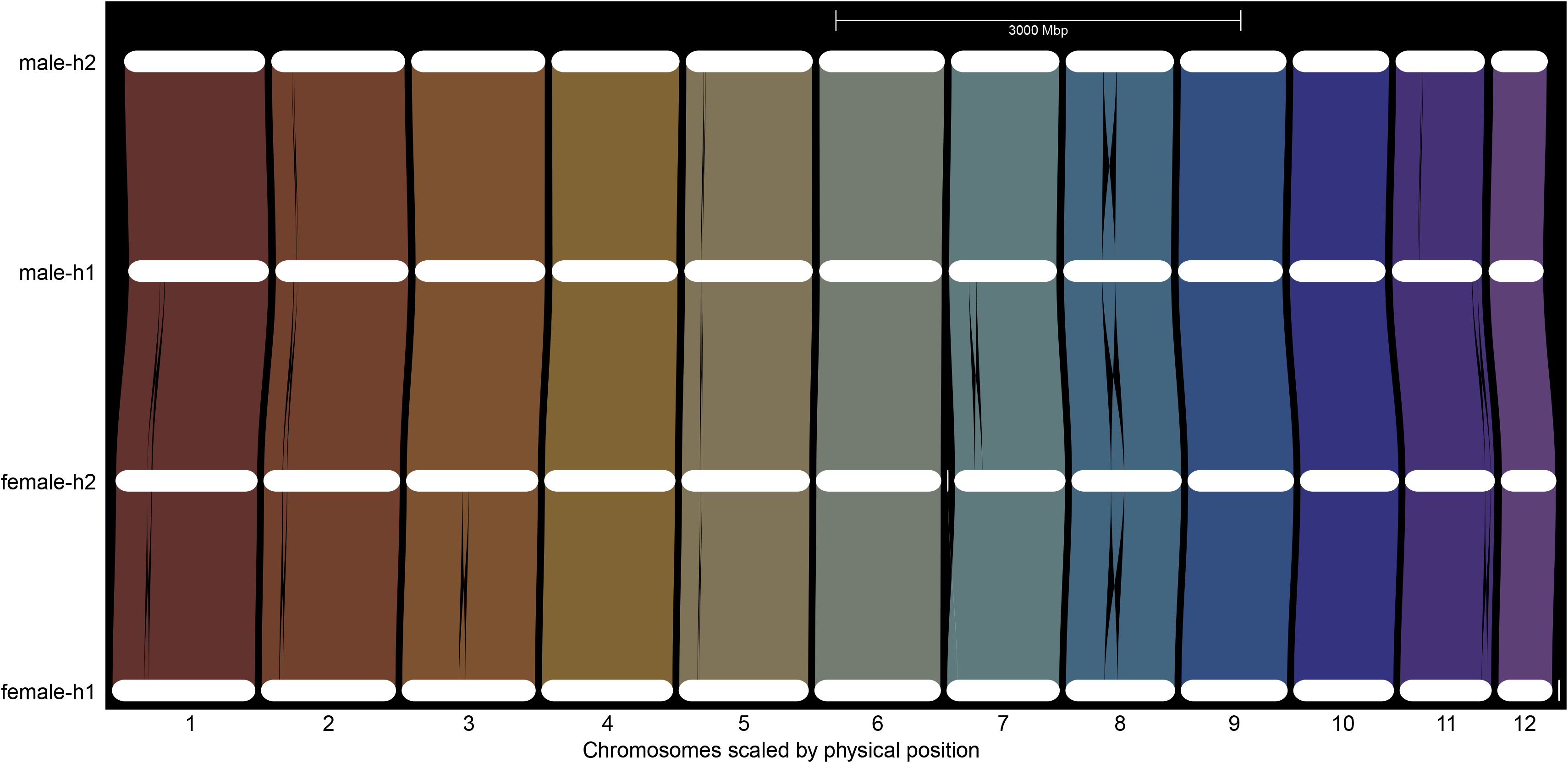
Riparian plot of synteny between the four assemblies based on orthologs in the structural annotation with GENESPACE shows similar results to the alignment-based synteny analysis with SyRI in Extended Data Fig. 2. The four assemblies are arranged one below the other in the following order: Tb_male-h2, Tb_male-h1, Tb_female-h2, Tb_female-h1. The 12 chromosomes are displayed separately, with additional small genomic segments in Tb_female-h2 after chromosome 6 and Tb_female-h1 after chromosome 12, both of which show no syntenic relationship with any other assembly. Syntenic relationships between adjacent assemblies are displayed as colored areas, with one color per chromosome. Inversions are displayed as hourglass-shaped blocks.

**Extended Data Figure 4.**
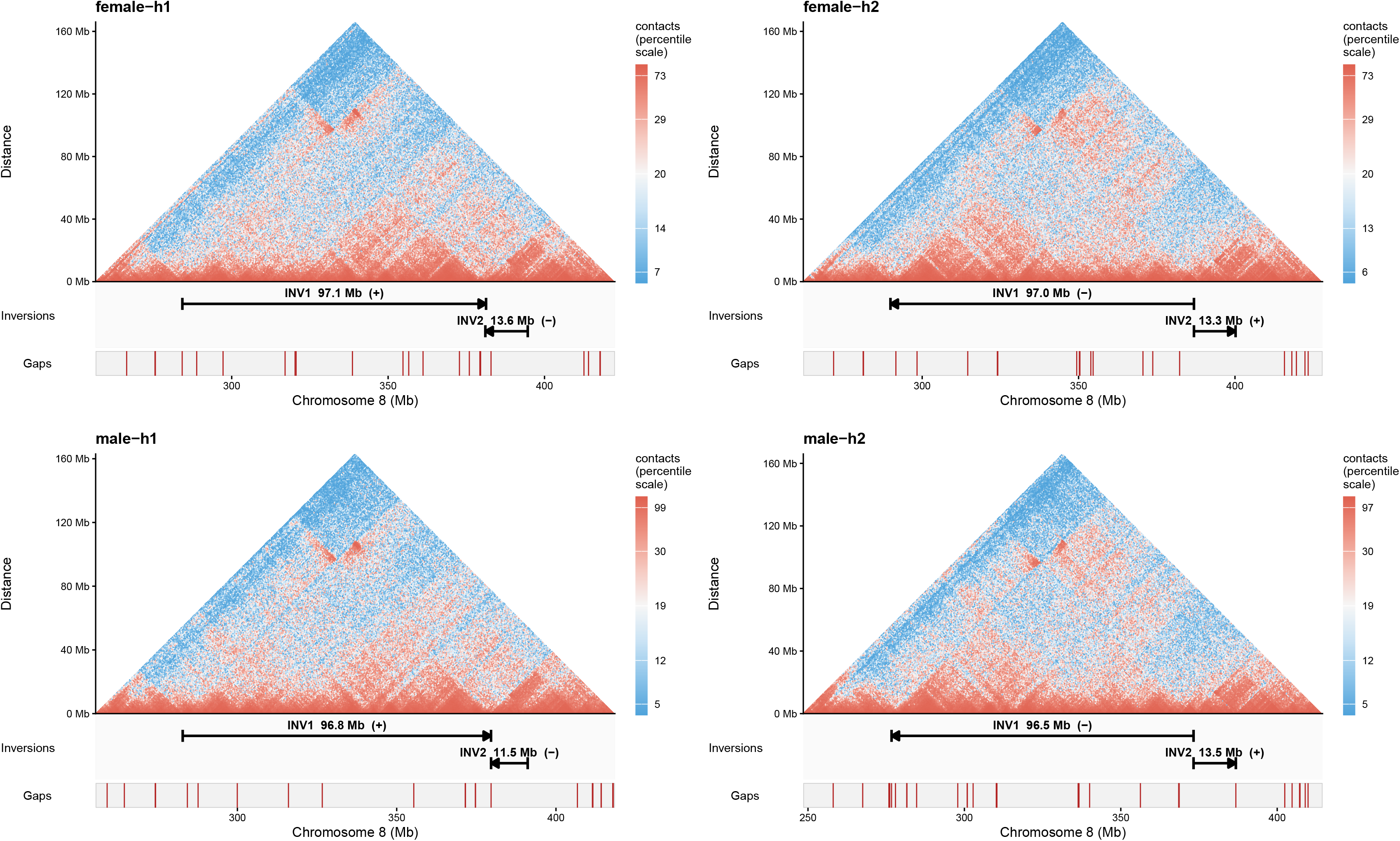
Highly resolved Hi-C contact heatmaps confirm the largest structural variant identified, a ∼97 Mb-inversion on chromosome 8 heterozygous in both individuals. Hi-C contact maps support the ∼97 Mb and ∼13 Mb inversions on chromosome 8 (INV1 and INV2; Tb_female-h1: 284.2–381.3 Mb and 381.3–394.6 Mb), heterozygous in both individuals. They are located directly adjacent to each other. Hi-C contact maps of chromosome 8 are presented for Tb_female-h1 (upper left panel), Tb_female-h2 (upper right panel), Tb_male-h1 (lower left panel) and Tb_male-h2 (lower right panel). For each panel, all Hi-C reads of the respective individual were mapped to that single haplotype assembly, so that reads from both homologous chromosomes are shown on the same reference. Contacts were binned at 10 kb, aggregated to multiple resolutions with cooler and are displayed as unnormalized counts at 500 kb resolution. The inversions are displayed by arrowheads and their directions compared to their complements are indicated by (+) and (-) labels. In all four assemblies, the main diagonal along the bottom of the heatmap shows a consistent signal of high interaction (red) compared to regions with medium or low interaction (white or blue, respectively). The continuous main diagonal (reads of the co-linear homolog) is accompanied by an off-diagonal “bowtie” signal (reads of the homolog carrying the segments in opposite orientation) in all four haplotypes, as expected for an inversion heterozygous in both individuals. The track below each map shows assembly gaps; none coincides with a breakpoint (see also Supplementary Fig. 2).

**Extended Data Figure 5.**
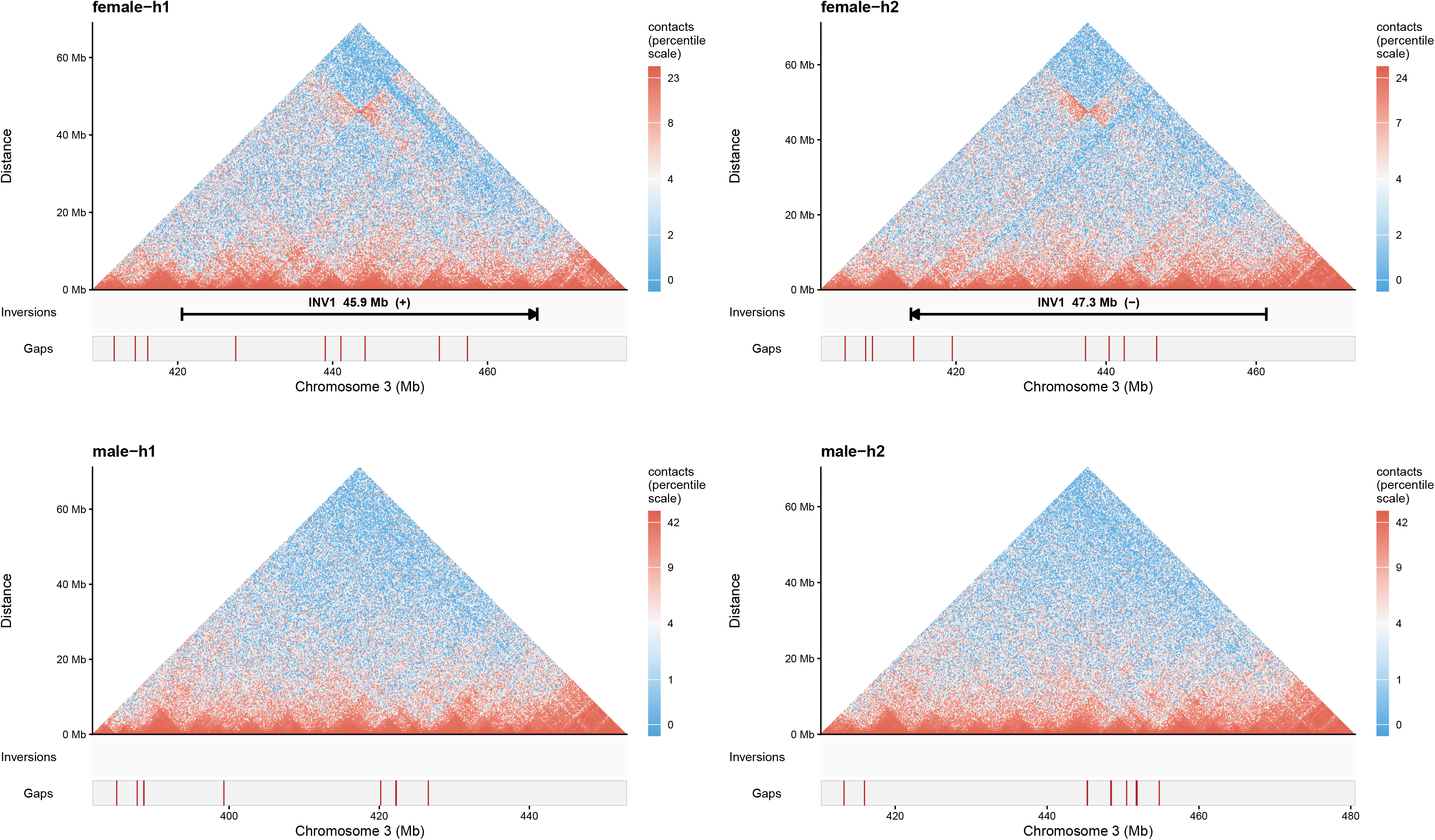
Highly resolved Hi-C contact heatmaps support the largest structural variant that is heterozygous only in the female individual, a ∼46 Mb-inversion on chromosome 3. Hi-C contact maps support an inversion on chromosome 3 (∼46 Mb) which is heterozygous only in the female (INV1; Tb_female-h1: 420.5–466.4 Mb). The maps were generated and displayed as in Extended Data Fig. 4: Tb_female-h1 (upper left panel), Tb_female-h2 (upper right panel), Tb_male-h1 (lower left panel) and Tb_male-h2 (lower right panel). Contacts were binned at 10 kb, aggregated to multiple resolutions with cooler and are displayed as unnormalized counts at 200 kb resolution. In each of the four assemblies, the main diagonal along the bottom of the heatmap shows a consistent signal of high interaction (red) compared to regions with medium or low interaction (white or blue, respectively). Female maps show the main diagonal plus the off-diagonal “bowtie” (heterozygous inversion); male maps show only the continuous diagonal (homozygous), therefore no inversion is marked. Gap tracks as in Extended Data Fig. 4. No gaps lie at or near the borders of an inversion.

**Extended Data Figure 6.**
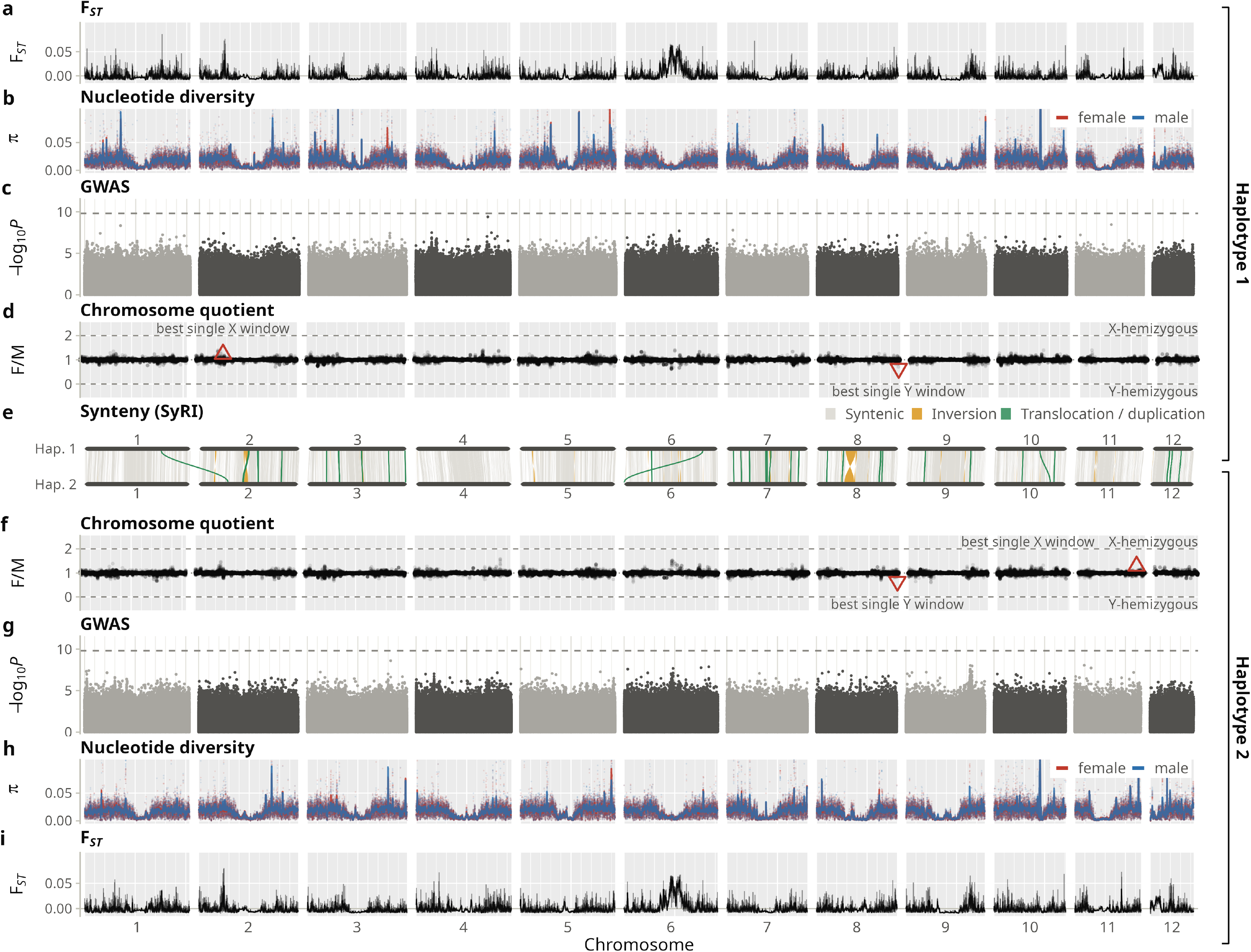
Reference-based analyses of 100 individuals across the two male haplotypes fail to identify a sex-determining region in *T. baccata*. **a,i** Genetic differentiation (*F_ST_*) between females and males was calculated in 100 bp windows and visualized as 100 kb averages. The black line shows the average across each plotting bin of about 2.4 Mb, the shaded band the range. **b,h** Nucleotide diversity π for females (red) and males (blue) in 100 kb windows is shown as points, with solid blue and red lines marking a sliding average across 5 Mb regions. **c,g** Manhattan plots show GWAS results (i.e., -log10 *P*-values) for sex of ca. 321 million biallelic SNPs. Horizontal dashed lines indicate 5% Bonferroni significance cutoff. **d,f** The chromosome quotient was calculated as the ratio of mean female to mean male (F/M) short-read DNA sequencing coverage in blocks of 1,000 consecutive 100 bp windows (∼100 kb), normalized by the genome-wide median depth of each sample. Only windows with a normalized depth of at least 0.2 in at least 25 of the 50 males were kept. Single best X (ratio closest to X-hemizygosity of 2) and Y (ratio closest to Y-hemizygosity of 0) 100 bp windows are shown as red triangles. **e** SyRI was used to assess syntenic regions (gray), inversions (orange) and translocations/duplications (green) across the 12 chromosomes. The upper half of the figure depicts male haplotype 1 (Tb_male-h1), the lower half male haplotype 2 (Tb_male-h2), which represent the two haplotypes of the male reference assemblies. The 12 chromosomes are shown on the x-axis. All analyses used 100 resequenced individuals, split into females (n=50) and males (n=50).

