## Supplementary Figures for "Haplotype-resolved genome assemblies and population resequencing reveal no evidence for genetic sex determination in the dioecious conifer Taxus baccata"

**This file includes:**

Supplementary Figures 1-12

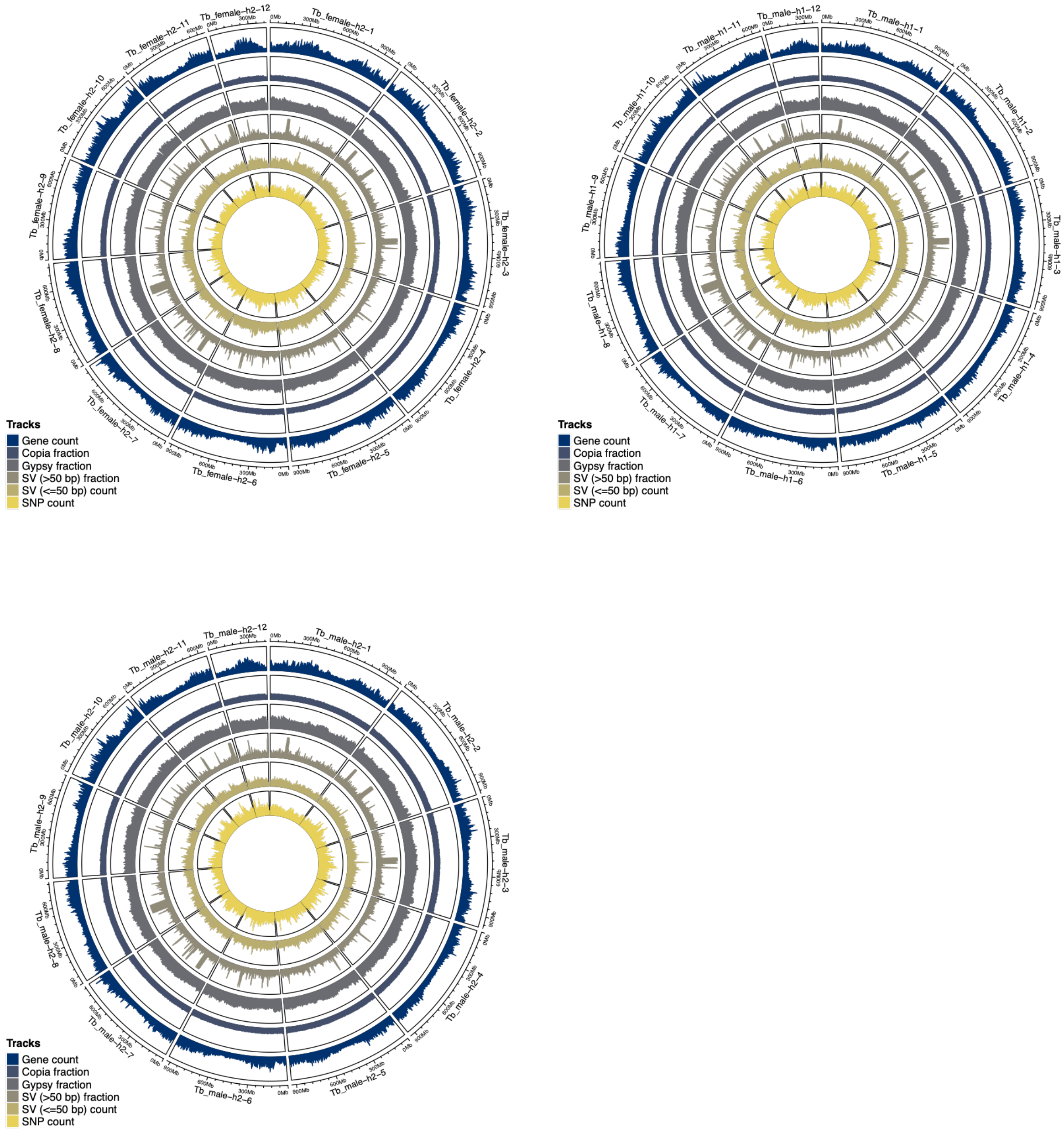


**Supplementary Figure 1: Chromosome-level assemblies for female haplotype 2, male haplotype 1 and male haplotype 2.** Circos plots of the Tb_female-h2, Tb_male-h1 and Tb_male-h2 assemblies. Statistics were calculated in 5 Mb sliding windows with 2.5 Mb steps. The following feature types are presented: Genes (counts per window); Copia elements, Gypsy elements, and large structural variants (> 50 bp) (fraction of the window covered by the respective feature type); small structural variants (<= 50 bp) and SNPs (counts per window).

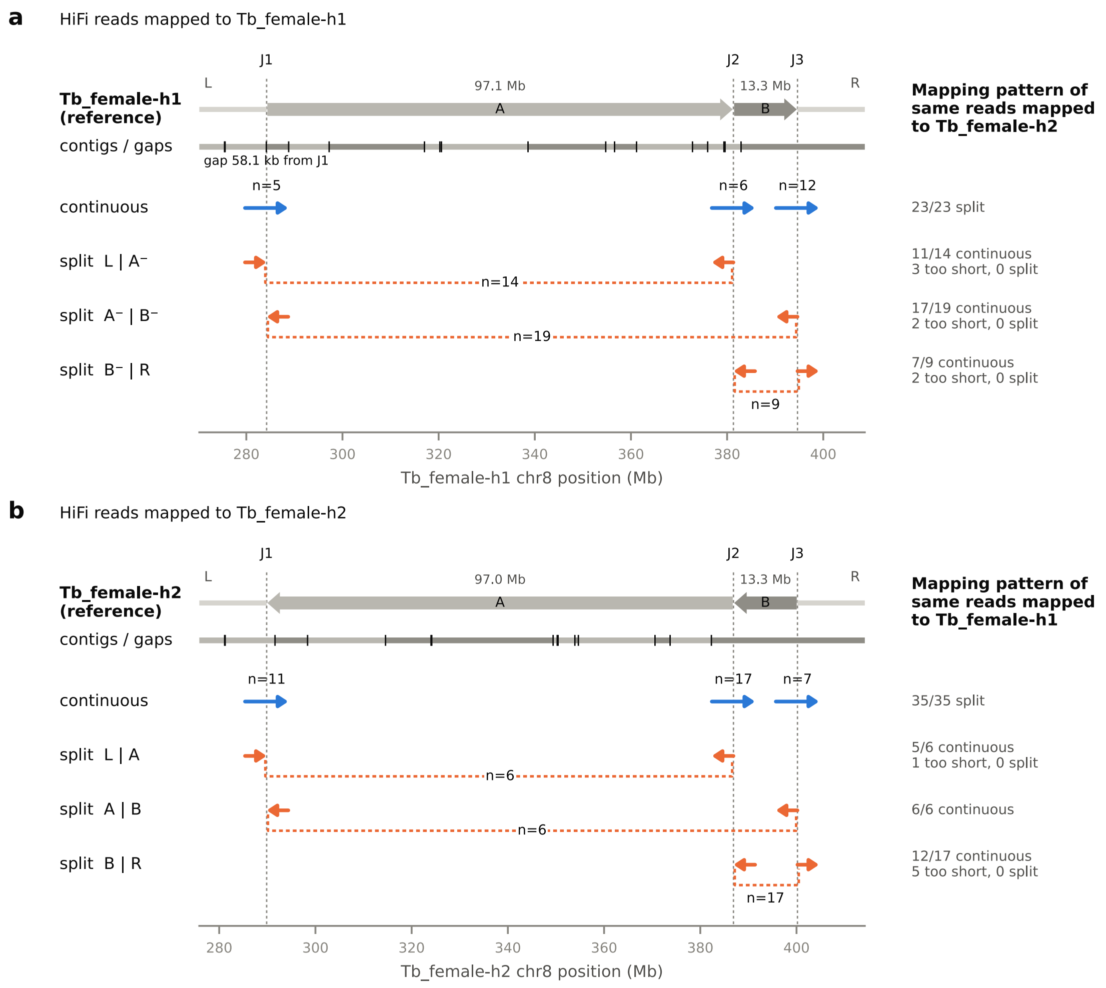


**Supplementary Figure 2. PacBio HiFi long-read mappings provide evidence for the large chromosome 8 inversions.** (**a, b**) HiFi reads of the female were aligned separately to each of its haplotype assemblies, i.e. Tb_female-h1 (a) and Tb_female-h2 (b). The top track of each panel shows the reference haplotype, with the two adjacent inverted segments (A = 97.1/97.0 Mb and B = 13.3 Mb) as grey arrows, oriented relative to Tb_female-h1. The flanks (L and R) are marked by grey lines, the three junctions (J1–J3) by vertical dotted lines. The track below shows the contigs of the chromosome-scale scaffold in alternating grey color, separated by assembly gaps indicated by black tick marks. None of the junctions coincides with a gap. The closest gap is 58.1 kb downstream from J1 in Tb_female-h1. Blue and orange arrows show continuous (with at least 1 kb on each side) and split long-read alignments, respectively. Split positions in reads perfectly agree (1–25 bp) with the breakpoints predicted by the assemblies. The number of reads with MAPQ ≥ 20 is indicated above the arrows. The right column classifies the same reads when mapped to the other haplotype assembly. All reads that are continuous when mapped to one assembly are split when mapped to the other. Reads that align across the matching junction with less than 1 kb on either side are classified as too short.

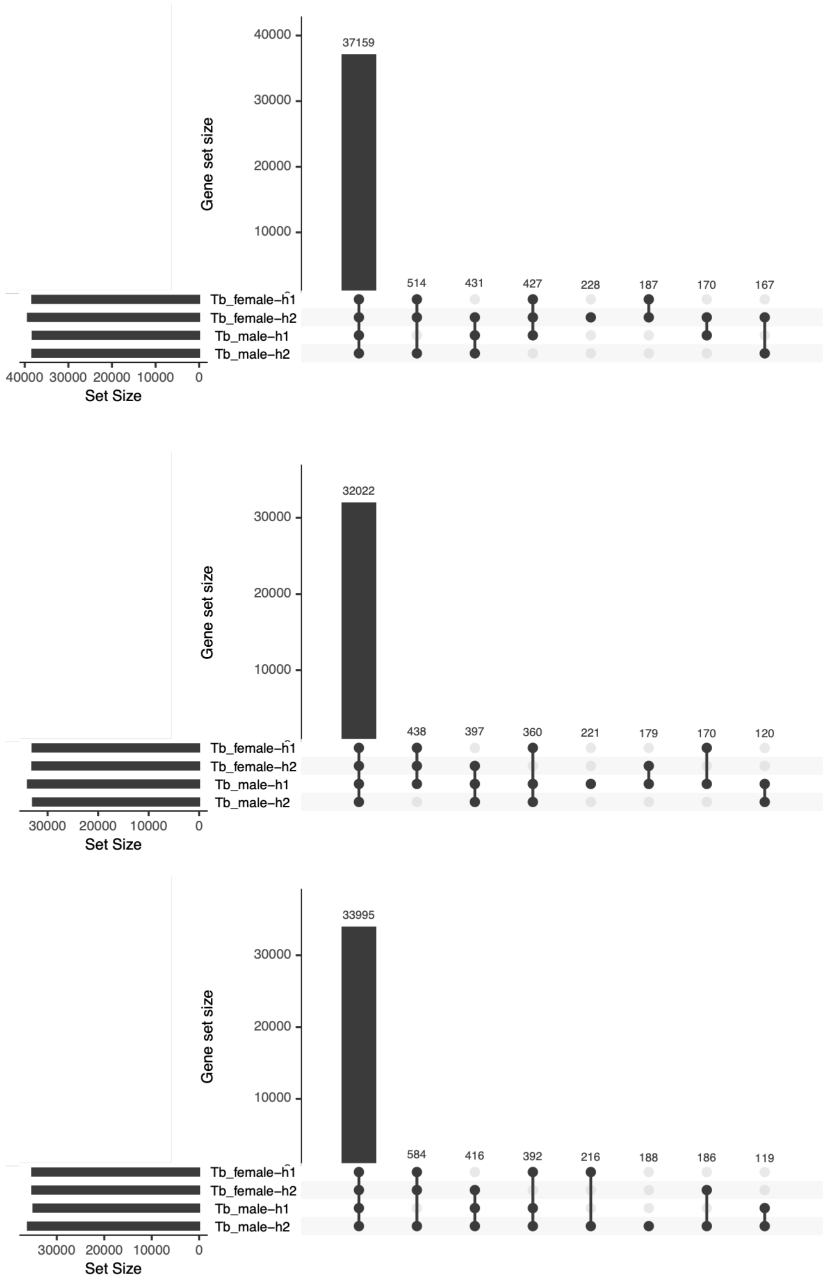


**Supplementary Figure 3: The assemblies and annotations of female haplotype 2, male haplotype 1 and male haplotype 2 show gene presence-absence variation.** Comparisons of the gene sequences annotated in the Tb_female-h2, Tb_male-h1 and Tb_male-h2 assemblies with the other assemblies' sequences, as determined by Liftoff, reveal presence-absence variation of genes. Each bar refers to one combination indicated below the bar, with black dots marking presence and gray dots absence from the assembly.

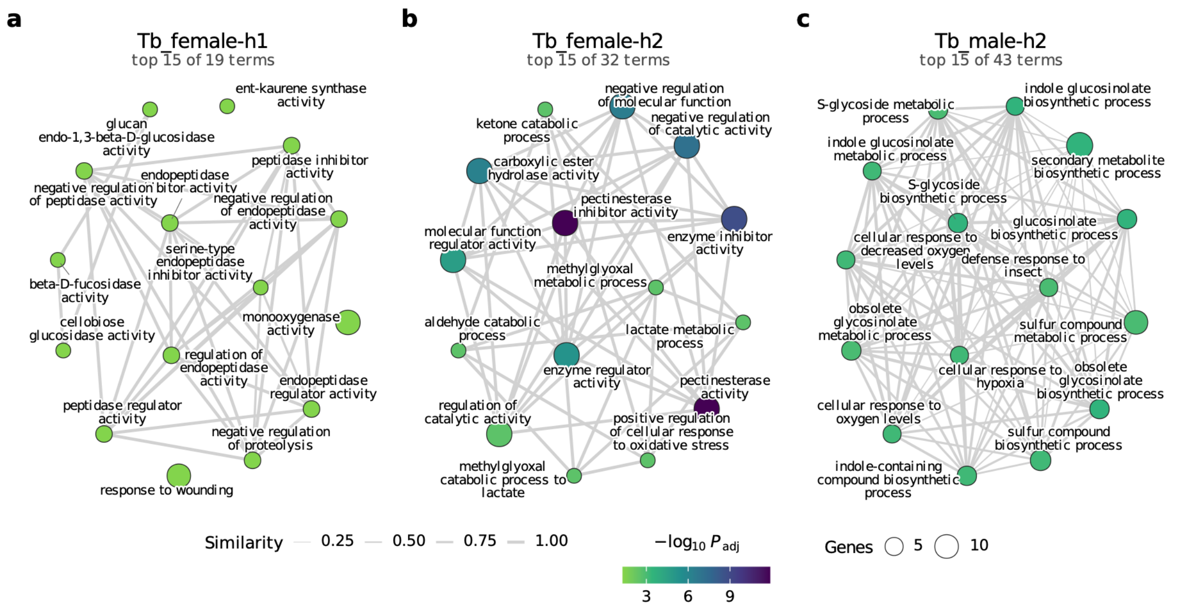


**Supplementary Figure 4: Enrichment maps of Gene Ontology (GO) terms overrepresented among haplotype-specific genes. a**, Tb_female-h1, 123 GO-annotated haplotype-specific genes, 19 significant terms. **b**, Tb_female-h2, 96 GO-annotated haplotype-specific genes, 32 significant terms. **c**, Tb_male-h2, 79 GO-annotated haplotype-specific genes, 43 significant terms. Terms with an adjusted *p*-value < 0.05 were considered significant. Each panel shows the 15 terms with the lowest adjusted *p*-values. No term was significant for the assembly Tb_male-h1 (for the 91 GO-annotated haplotype-specific genes). Every node represents a GO term. Node size corresponds to the number of haplotype-specific genes annotated to the term and color represents -log10 $P_{adj}$. Edges connect two terms whose haplotype-specific genes overlap with a Jaccard index >= 0.2. Line width indicates a larger overlap (i.e., higher "similarity"). Color, size and width scales are shared across panels.

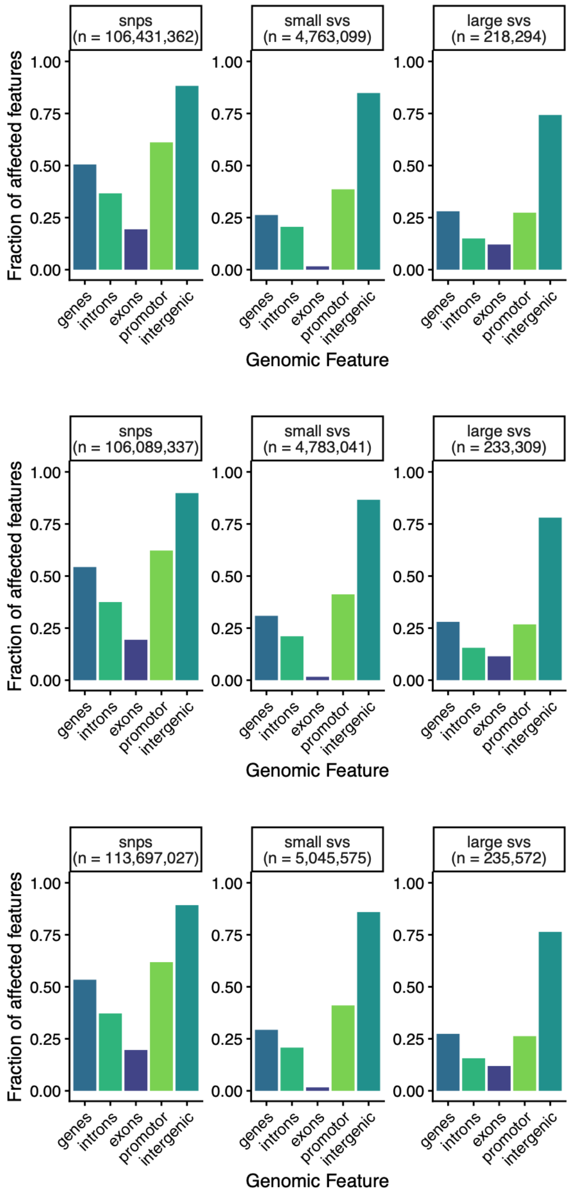


**Supplementary Figure 5: The assemblies of female haplotype 2, male haplotype 1 and male haplotype 2 show high levels of sequence variation between haplotypes and conservation of genes.** Fractions of genomic features that overlap with different types of variants, i.e., SNPs, and small (<= 50 bp) or large structural variants (> 50 bp) reported by SyRI (Supplementary Table 2) show conservation of genes, and specifically exons compared to intergenic regions. The regions up to 5 kb upstream of a gene are considered promoter regions.

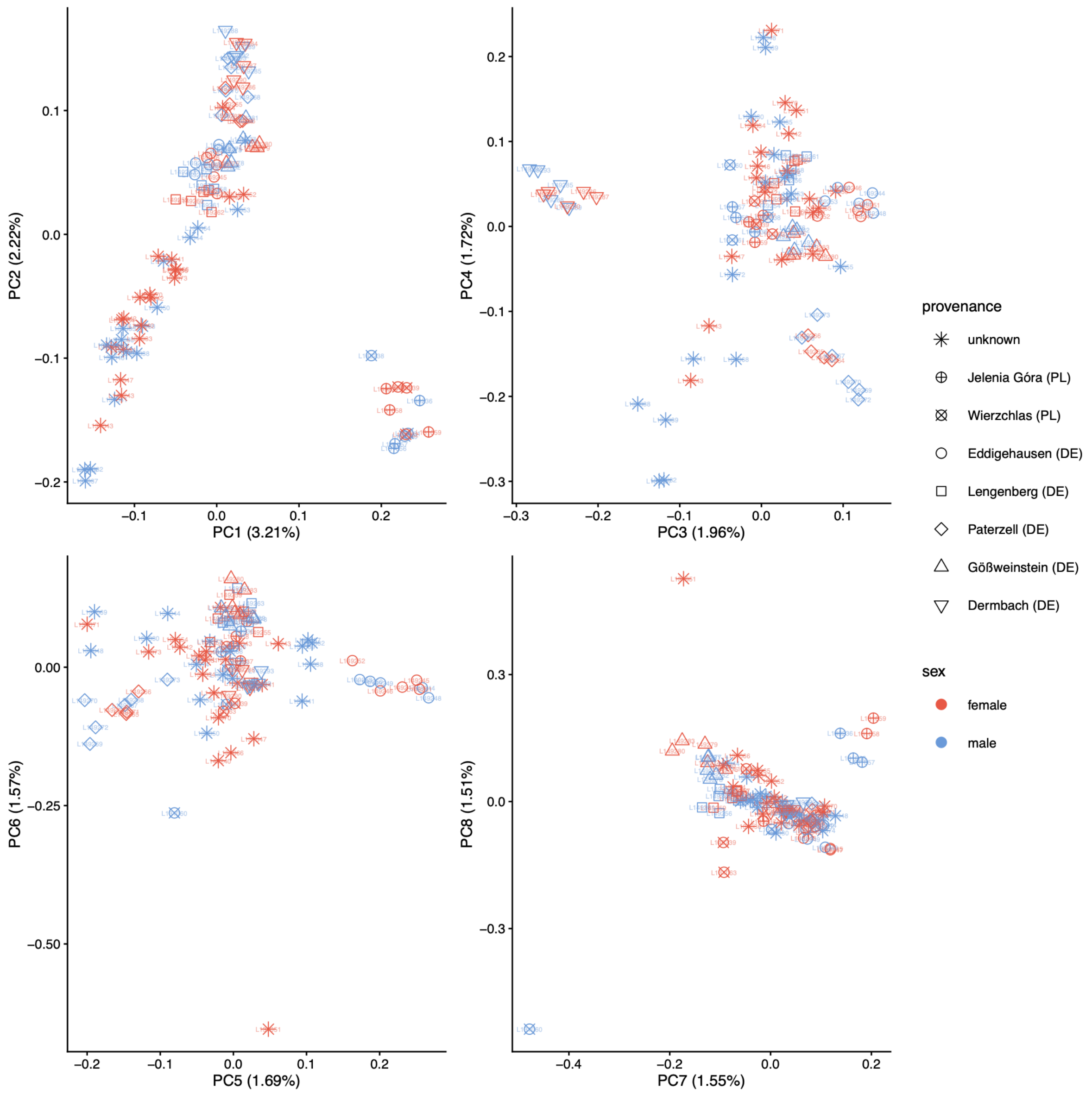


**Supplementary Figure 6: Genomic variation shows population structure in *Taxus baccata* across the sampling locations.** PCA based on 21M largely unlinked genome-wide SNPs called from mappings to Tb_female-h1 shows population structure among the 50 female (red) and 50 male (blue) *T. baccata* individuals. Each point depicts one individual. Shapes indicate origins.

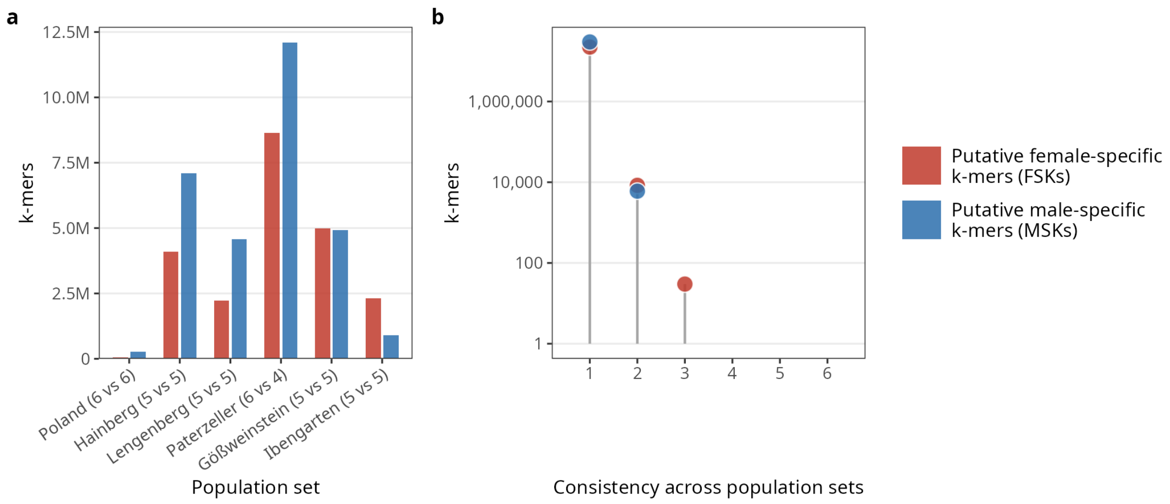


**Supplementary Figure 7: Reference-free k-mer analyses in *T. baccata* natural populations provide no evidence of sex-specific sequences. a,** Counts of putative sex-specific k-mers in 6 natural populations with 5 to 6 female and 4 to 6 male samples. Sets are ordered based on their FSK/MSK ratio. **b,** Consistency of FSKs and MSKs across the sets shown in a. Consistency refers to the number of sets that contain a given FSK or MSK.

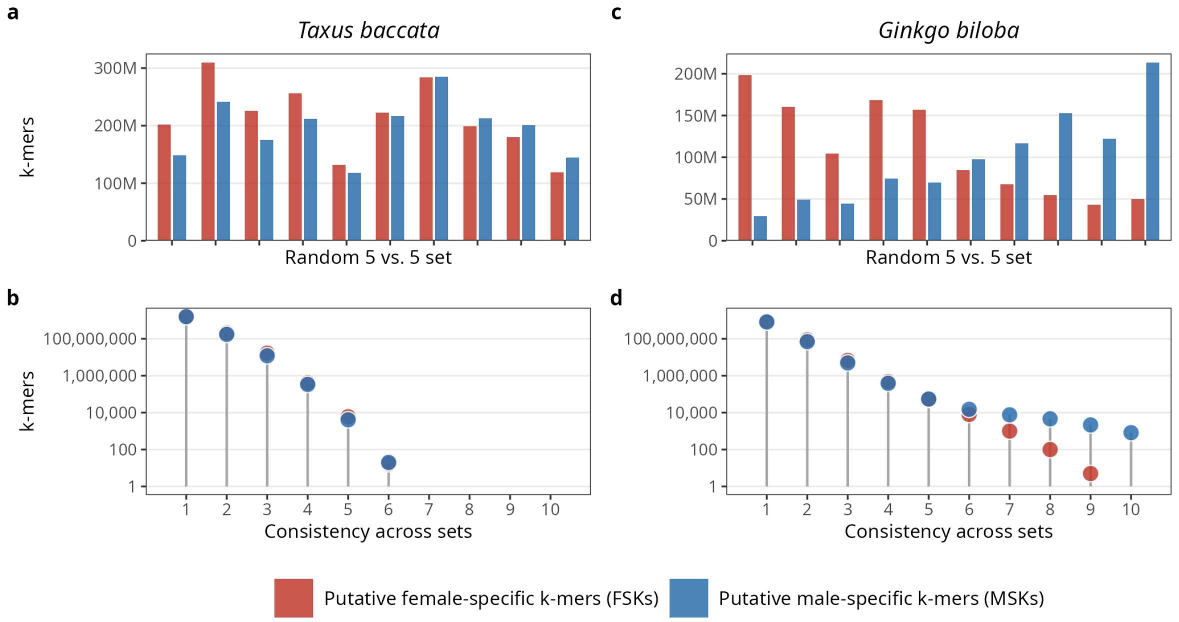


**Supplementary Figure 8: Reference-free k-mer analyses in *T. baccata* and *Ginkgo biloba* provide no evidence of sex-specific sequences in *T. baccata* even when allowing one outlier in each group. a,c** Counts of putative sex-specific k-mers in 10 random sets of 5 female and 5 male samples of *T. baccata* (a) and *G. biloba* (c) allowing one outlier per group (i.e., among females and males) demonstrate a high degree of noise in both species. Sets are ordered based on their FSK/MSK ratio. **b,d** Consistency of FSKs and MSKs across the sets shown in a,c. Consistency refers to the number of sets that contain a given FSK or MSK.

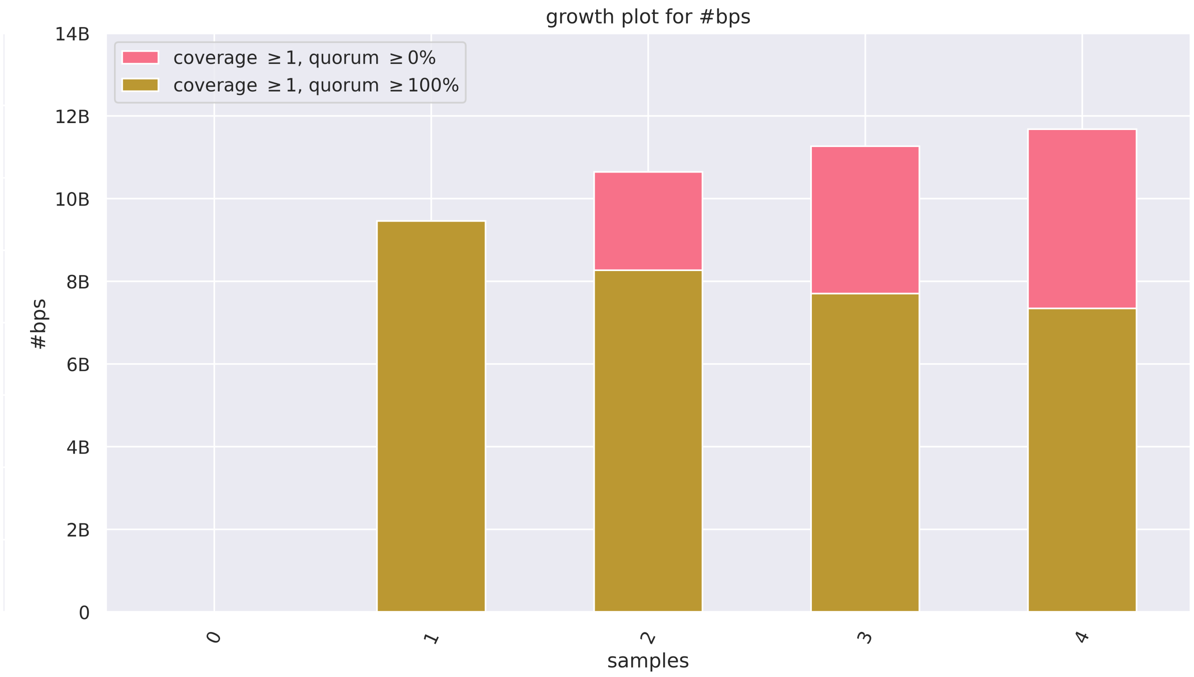


**Supplementary Figure 9: Growth plot of a *T. baccata* pangenome graph.** The y axis shows the number of non-redundant base pairs in the Minigraph-Cactus graph and the x axis the number of haplotypes added. Values are averaged over all possible orders of adding haplotypes. The total bar height is the pangenome size. Dark yellow is the core sequence shared by all added haplotypes. The pink part above the core is the dispensable (variable) sequence not shared by all haplotypes. Quorum is the minimum fraction of haplotypes that must traverse a base pair for it to be counted.

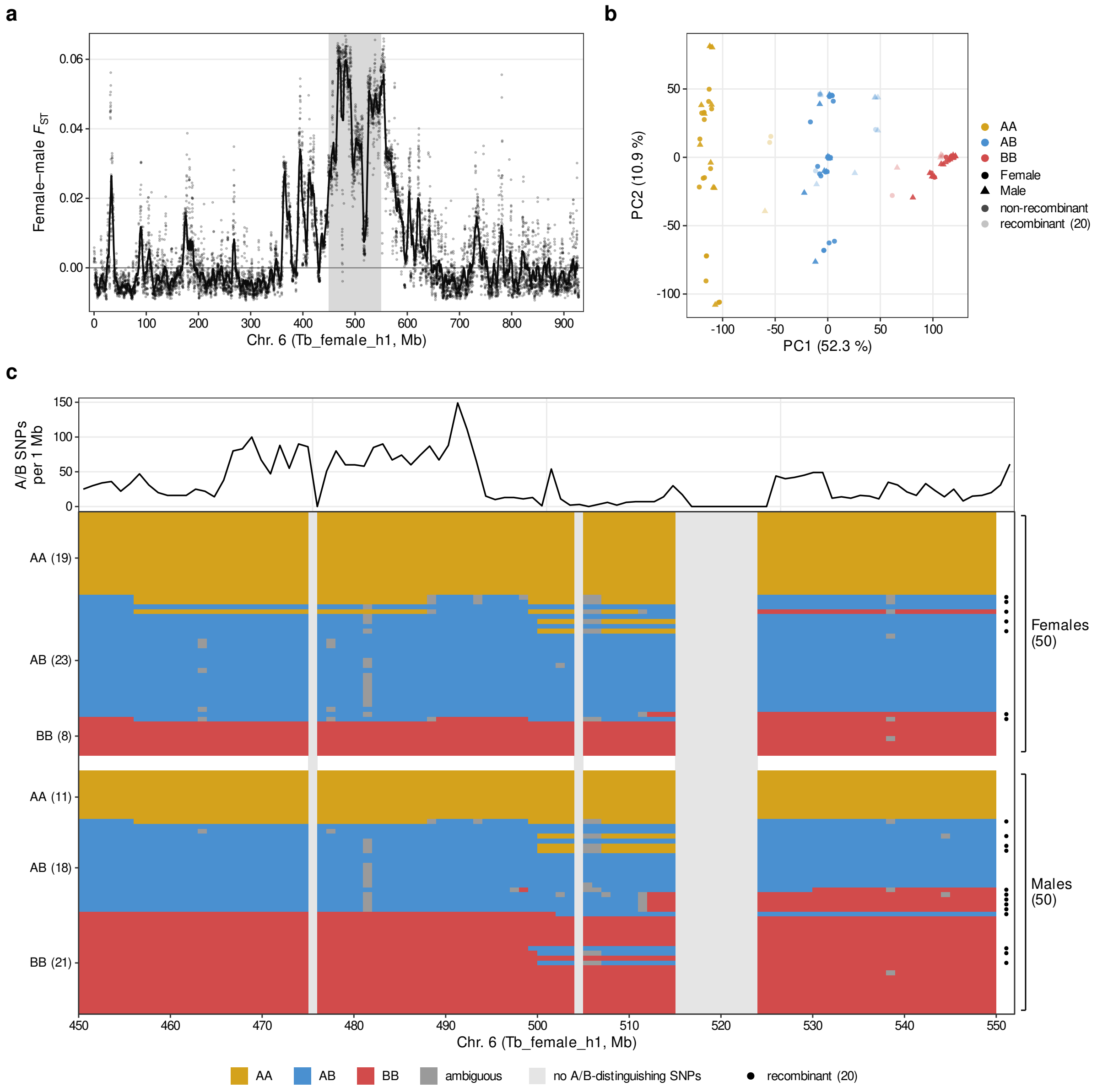


**Supplementary Figure 10. A large haplotype block on chromosome 6 exhibits increased genetic differentiation between the sexes but is not consistent with a sex-determining region.** (**a**) Genetic differentiation (Hudson's $F_{ST}$) between 50 female and 50 male *T. baccata* individuals is shown along chromosome 6 of Tb_female_h1 with 100 kb averaged windows depicted as points. The line indicates a rolling mean across 5 Mb windows. The shaded area from 450-550 Mb is analyzed in detail in panels b and c. (**b**) PCA using 13,135 SNPs (every 100$^{th}$ biallelic SNP with minor allele frequency > 0.1 and missingness < 10% in the region) shows three separate clusters on PC1 corresponding to two haplotype blocks, which are either homozygous (AA = orange, BB = red) or heterozygous (AB = blue). Trees were assigned to the three clusters (AA, AB, BB) by k-means clustering of their PC1 values. Circles depict females, triangles males. Recombinant individuals (with more than 10% of their genotypes deviating from their main cluster) are marked by transparent color. (**c**) The 3,390 SNPs that exhibit allele frequency differences of more than 0.9 between AA and BB trees ("A/B SNPs") were used to analyze the genotypes in 1 Mb bins across the two haplotype blocks in the females (top 50 rows) and males (bottom 50 rows). Genotypes for each bin were assigned by B allele dosage (0-0.25=AA, 0.75-1.25=AB, 1.75-2=BB). Light-grey columns indicate bins without any SNPs distinguishing the A and B haplotypes. The number of A/B SNPs is shown on top. Dots on the right mark the 20 recombinant trees.

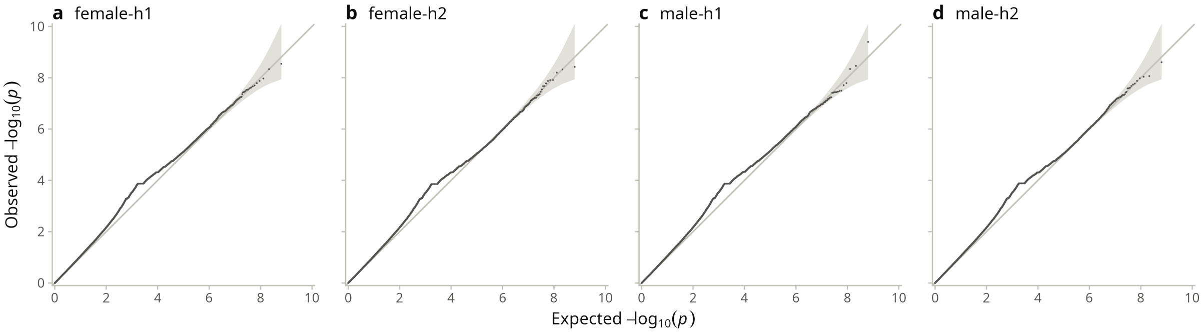


**Supplementary Figure 11: QQ-plots of the GWAS for sex against the four genome assemblies.** Expected vs. observed -log10(*p*) values for the GWAS against the two female (a,b), n=320M each, and two male (c,d), n=321M each, genome assemblies. The diagonal line marks a perfect 1:1 correlation. The shaded area indicates the pointwise 95% confidence interval. The bump around 4 corresponds to the randomly linked variants of the chromosome 6 haplotype block (Supplementary Fig. 10).

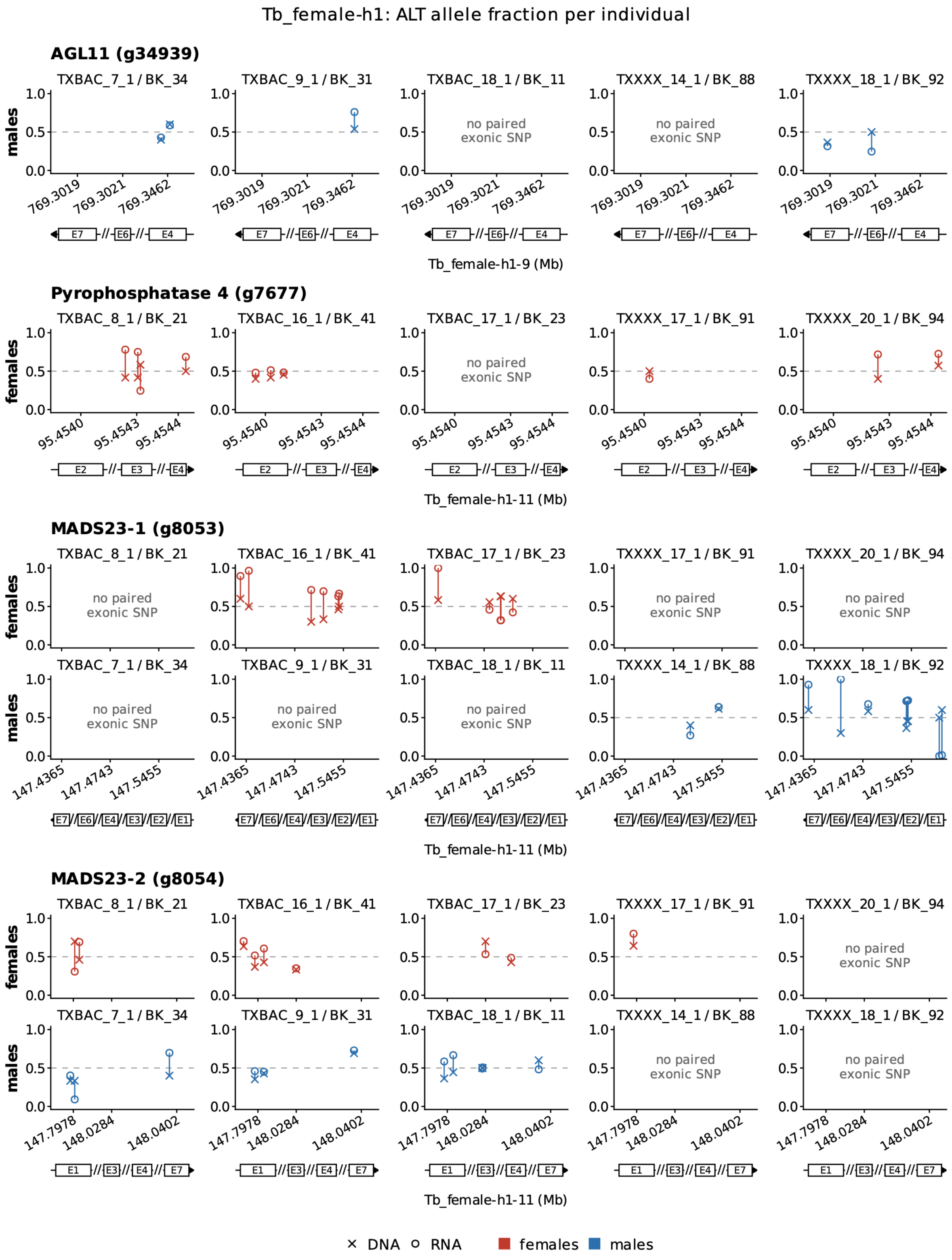


**Supplementary Figure 12: Alternative (ALT) allele fraction per individual shows biallelic expression of candidate sex-determining genes.** Mean fraction of female and male individuals of the alternative allele (ALT) at selected heterozygous SNP positions (chromosomal positions in exons, marked by E and the exon number) called in short read data from DNA sequencing (DNA) or RNA sequencing (RNA) mapped against female-h1.
